# Region Capture Micro-C reveals coalescence of enhancers and promoters into nested microcompartments

**DOI:** 10.1101/2022.07.12.499637

**Authors:** Viraat Y. Goel, Miles K. Huseyin, Anders S. Hansen

## Abstract

Although enhancers are central to the regulation of mammalian gene expression, the mechanisms underlying Enhancer-Promoter (E-P) interactions remain unclear. Chromosome conformation capture (3C) methods effectively capture large-scale 3D genome structure but struggle to achieve the depth necessary to resolve fine-scale E-P interactions. Here, we develop Region Capture Micro-C (RCMC) by combining MNase-based 3C with a tiling region-capture approach and generate the deepest 3D genome maps reported thus far with only modest sequencing. By applying RCMC in mouse embryonic stem cells and reaching the genome-wide equivalent of ∼200 billion unique contacts, RCMC reveals previously unresolvable patterns of highly nested and focal 3D interactions, which we term microcompartments. Microcompartments frequently connect enhancers and promoters and are largely robust to loss of loop extrusion and inhibition of transcription. We therefore propose that many E-P interactions form through a compartmentalization mechanism, which may explain why acute cohesin depletion only modestly affects global gene expression.

## INTRODUCTION

3D genome structure regulates vital cellular processes including gene expression, DNA repair, genome integrity, DNA replication, and somatic recombination^1,2^. Many insights into 3D genome structure have come from Chromosome Conformation Capture (3C) assays, which have revealed structural hallmarks across at least three scales. First, active and inactive chromatin segregate into A- and B-compartments through a poorly understood compartmentalization mechanism^3,4^. Second, the genome is folded into loops and local domains called Topologically Associating Domains (TADs) or loop domains^5–8^ by loop-extruding cohesin complexes halted at CTCF boundaries^9,10^. Third, while A/B-compartments and TADs generally span hundreds to thousands of kilobases, recent work has hinted at finer scale 3D chromatin interactions including between enhancers and promoters^11–17^. Because enhancers are the primary units of gene expression control in mammals, there has been intense interest in resolving fine-scale enhancer-promoter (E-P) interactions. However, it has remained challenging to resolve fine-scale E-P interactions with current methods^8,18^. This motivated us to develop a new 3C method that effectively captures E-P interactions.

Advances in our understanding of 3D genome structure have been primarily driven by: (1) deeper sequencing; (2) improved 3C protocols; and (3) perturbation studies. First, A/B-compartments, TADs, and loops were uncovered as deeper sequencing increased the number of captured unique contacts in 3C experiments from ∼8 million^3^ to ∼450 million^5^ to ∼5 billion^7^, respectively. Second, in overcoming the resolution limits imposed by Hi-C’s dependence on restriction enzymes, Micro-C achieved nucleosome-scale resolution by digesting chromatin with micrococcal nuclease (MNase); this allows Micro-C to better resolve finer-scale regulatory interactions including between enhancers and promoters^8,11–13,15,19,20^. Third, perturbation studies have yielded profound mechanistic insights into 3D genome structure. For example, protein-depletion studies were pivotal in elucidating the roles of CTCF, cohesin, and associated factors in the formation of TADs and loops^12,21–27^.

Nevertheless, despite decreasing sequencing costs, sequencing remains the key bottleneck for 3C assays. For a genome with *n* bins, sequencing costs to populate an *n*^2^ pairwise contact matrix grow quadratically with *n*. For example, we estimate approximately $1.6 billion in sequencing costs alone to average one read per nucleosome-sized bin across the human genome (a total of (3.3×10^9^ bp/150 bp)^2^/2 = 2.4×10^14^ reads). To overcome the prohibitive cost of sequencing inherent to current methods and facilitate the study of fine-scale 3D genome structure and enhancer-promoter interactions, we therefore sought to develop a 3C method that (1) strongly increases effective sequencing depth, (2) incorporates the latest advances in 3C-derived protocols, and (3) is cost-effective for perturbation experiments.

Here, we address these three points by combining Micro-C with a tiling region capture approach^28,29^ to enrich for entire regions of interest in a new method we call Region Capture Micro-C (RCMC). We use RCMC to generate the deepest maps of 3D genome organization reported so far, achieving nucleosome resolution with a fraction of the sequencing. By reaching the local equivalent of ∼200 billion unique contacts genome-wide, we discover patterns of previously unseen, fine-scale, focal, and highly nested 3D interactions in gene-dense loci that we call microcompartments. Microcompartments frequently connect enhancers and promoters, and require neither loop extrusion nor transcription. Taken together, our results suggest that interactions between enhancers and promoters, now highly resolved by RCMC, may be driven by compartmentalization mechanisms rather than loop extrusion.

## RESULTS

### Region Capture Micro-C (RCMC): Development, validation, and benchmarking

To develop Region Capture Micro-C (RCMC), we optimized the regular Micro-C protocol^11,13,15^ to maximize library complexity and combined it with a tiling region capture approach^28,29^ (**Fig. 1a**). Briefly, mouse embryonic stem cells (mESCs) were crosslinked with disuccinimidyl glutarate (DSG) and formaldehyde (FA) and digested to nucleosomes with MNase (**Fig. S1a-b**), after which fragment ends were repaired with biotin-labeled nucleotides and then proximity ligated. After protein removal and reversal of crosslinks, we size-selected and pulled down ligated dinucleosomal fragments, and prepared a Micro-C sequencing library. Avoiding repetitive regions, we designed 80-mer biotinylated oligos tiling 5 regions of interest, each spanning between 425 kb and 1,900 kb (**Fig. S1c**), and pulled-down the tiled regions of interest with 35-49% efficiency in a single step (**Fig. S1d**). After paired-end sequencing and normalization^30^ (**Fig. S1e**), we obtained contact maps (**Fig. 1a**). To validate our RCMC contact maps, we compared them to high-resolution Hi-C^31^ and Micro-C^13^ for the same regions. Our RCMC data matched both Hi-C^31^ and Micro-C^13^ data at 2-kb resolution (**Fig. S1f**), was reproducible (**Fig. S1g**), and gave the expected contact frequency scaling (**Fig. S2a**). Thus, RCMC captures all information in target regions obtained in prior multi-billion contact studies^13,31^.

**Figure 1.**
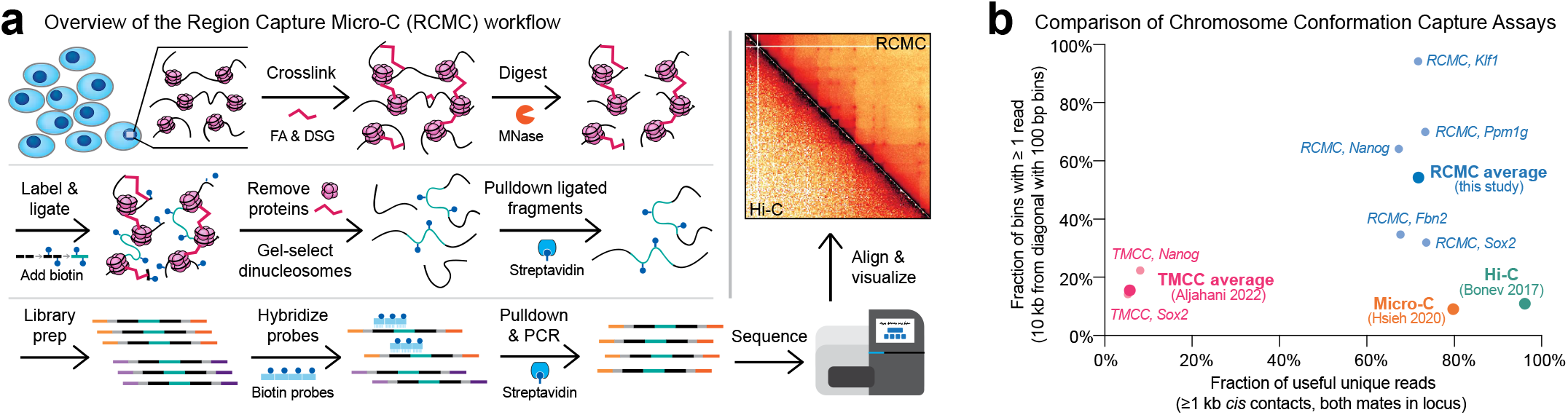
Region Capture Micro-C captures chromosome conformation at nucleosome resolution. (**a**) Overview of the Region Capture Micro-C (RCMC) protocol. Cells are chemically fixed, nuclei are digested with micrococcal nuclease (MNase), and fragments are biotinylated, proximity ligated, dinucleosomes gel extracted and purified, library prepped, PCR amplified, and region-enriched to create a sequencing library. After sequencing, mapping, and normalization, the data is visualized as a contact matrix. (**b**) Benchmarking comparison of RCMC against the highest resolution Tiled-Micro-Capture-C (TMCC)^17^, Micro-C^13^, and Hi-C^31^ mESC datasets. Region-averaged calculations are shown for RCMC, TMCC, Micro-C, and Hi-C, and calculations for individual captured regions are also shown for RCMC and TMCC. The x-axis shows the fraction of all reads that 1) uniquely map to the target region (both read mates fall within the Captured region) and 2) are structurally informative (*cis* contacts >=1 kb). The y-axis shows the fraction of all contact bins that contain at least one read using 100 bp bins 10 kb from the diagonal.

Having validated RCMC, we next benchmarked it against other 3C datasets. Despite capturing ∼2.6-3.3 billion unique contacts, the deepest Hi-C^31^ and Micro-C^13^ datasets in mESCs gave sparse contact maps (**Fig. 1b**). In contrast, since RCMC focuses its sequencing reads in only regions of interest, almost all interaction bins showed at least one interaction for our most deeply sequenced region (*Klf1* **Fig. 1b; Fig. S2b**). Indeed, with relatively modest sequencing (**Fig. S2c**) we captured the genome-wide equivalent of ∼200 billion unique contacts at the *Klf1* region.

To visualize the improvements afforded by RCMC, we plotted contact maps comparing RCMC to Hi-C^31^ and Micro-C^13^ at our 5 captured regions (**Fig. S3-4**). While A/B-compartments, TADs, and CTCF and cohesin-mediated structural loops are well-resolved in prior high-resolution Hi-C^31^ and Micro-C^13^ studies, resolving enhancer-promoter (E-P) interactions has proven more challenging^8,18^. To test the ability of RCMC to resolve E-P interactions, we captured a region around *Sox2* (**Fig. 2a**). *Sox2* encodes a key pluripotency transcription factor, whose expression in mESCs is controlled by a well-characterized ∼100 kb distal enhancer (Sox2 Control Region (SCR))^32–34^. While long-range *Sox2*-SCR interactions are visible in Hi-C and Micro-C, RCMC resolved the fine-scale substructure of the *Sox2*-SCR interactions: rather than one broad loop, *Sox2* forms multiple individual focal interactions with subelements of the SCR marked by Mediator binding and ATAC peaks (**Fig. 2a**). Furthermore, RCMC also revealed novel long-range interactions between a ∼600-700 kb distal region and *Sox2* and the SCR as well as strong compartmental exclusion of a ∼550 kb intervening region (**Fig. S4a**). Next, we focused on a ∼300 kb segment of our most deeply sequenced region, the region around *Klf1* (**Fig. 2b**). Notably, RCMC revealed patterns of highly focal and nested interactions in the *Klf1* region which are not visible in genome-wide Hi-C or Micro-C data (**Fig. 2b**). We name these interactions microcompartments (see Discussion for rationale and definition). We conclude that for specific regions, RCMC outperforms genome-wide Hi-C and Micro-C at a fraction of the cost.

**Figure 2.**
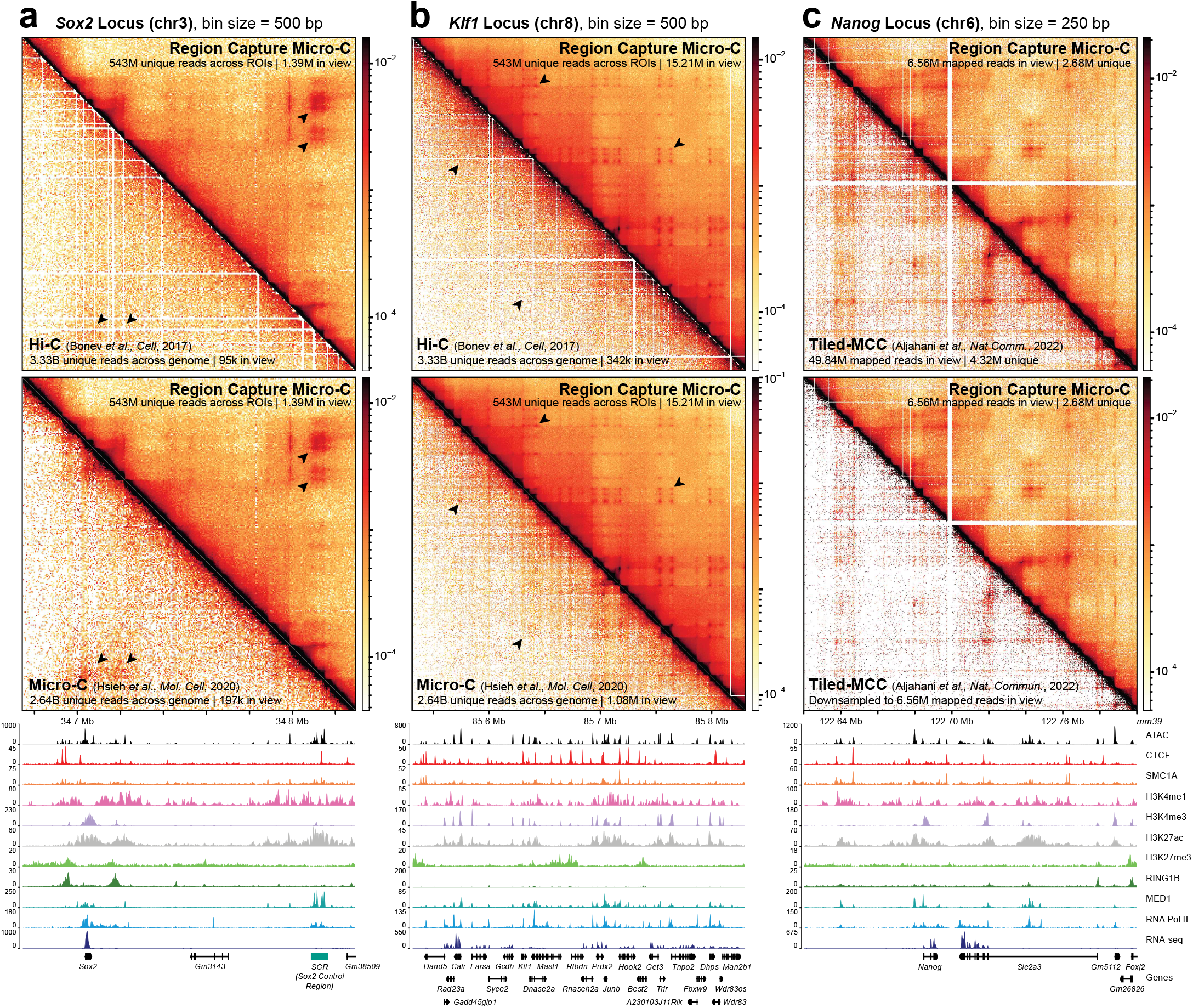
RCMC generates deep contact maps, reveals new aspects of 3D genome structure, and outperforms other 3C methods. (**a-b**) Contact map comparison of RCMC against the deepest available mESC Hi-C (top; Bonev 2017^31^) and Micro-C (middle; Hsieh 2020^13^) datasets at the (**a**) *Sox2* and (**b**) *Klf1* regions at 500 bp resolution. Gene annotations and ATAC, ChIP, and RNA-seq (see Supplementary Table 1) signal tracks are shown below the contact maps, while the contact intensity scale is shown to the right. The RCMC data shown throughout this manuscript were pooled from 3 replicates in wild-type mESCs. (**c**) Contact map comparison of RCMC against Tiled-Micro-Capture-C (TMCC)^17^ at the *Nanog* locus at 250 bp resolution. Full datasets are visualized in the top contact map, and TMCC has been downsampled to match the total number of RCMC sequencing reads in view in the bottom contact map.

Finally, while our studies were ongoing, the related methods Micro-Capture-C (MCC)^16^ and Tiled-Micro-Capture-C (TMCC)^17^ were reported. Unlike RCMC, (T)MCC uses only formaldehyde for fixation^35^, skips the pull-down of ligation products and the gel purification of dinucleosomes (**Fig. 1a**), and instead uses sonication to generate small fragments containing both ligated and unligated DNA. This allows (T)MCC to precisely sequence the ligation junction, which for RCMC would otherwise require longer-read sequencing. Thus, this affords (T)MCC base-pair resolution when capturing the interactions between regulatory elements^16,17^. However, by not enriching for the informative ligation products, (T)MCC mainly captures unligated DNA fragments, resulting in most sequencing reads being uninformative (**Fig. 1b**). Indeed, even with similar total sequencing reads, RCMC captured ∼134 million unique >1 kb *cis* contacts in the target regions compared to just ∼9-13 million for TMCC, underscoring the more than one order of magnitude higher efficiency of RCMC (**Fig. S2c)**. To directly compare RCMC to TMCC, we designed probes against the same *Nanog* region used in TMCC^17^. Due to the less efficient nature of TMCC, even with almost 10-fold higher sequencing at the *Nanog* region, TMCC maps were noisier than RCMC, which became even more evident when we subsampled TMCC to match RCMC (**Fig. 2c; Fig. S4b**). In summary, we conclude that RCMC is more efficient for general 3D genome structure mapping of a region, while (T)MCC may be applied when it is necessary to resolve ligation junctions with base-pair resolution.

### RCMC reveals highly nested and focal interactions between enhancers and promoters in gene-rich regions

RCMC data revealed highly nested and focal interactions in both the *Klf1* and *Ppm1g* regions which were not visible in multi-billion contact genome-wide Hi-C^31^ and Micro-C^13^ (**Fig. 2b, 3a-b, Fig. S5a-b**). We applied existing loop^36,37^ and compartment calling algorithms^37,38^ to identify these interactions, but they did not reliably detect them (**Fig. S5c**). We therefore manually identified 132 anchors forming a total of 1093 focal interactions in the gene-rich *Klf1* and *Ppm1g* regions (**Fig. 3a-b**; **Fig. S5d**). Furthermore, we validated that these interactions were not due to incomplete contact map normalization^30^ (**Fig. S6a**) nor an artifact of increased accessibility at the anchors (only about half of all ATAC peaks result in “dots” and not all “dots” are anchored by ATAC peaks; **Fig. S6b-d**).

**Figure 3.**
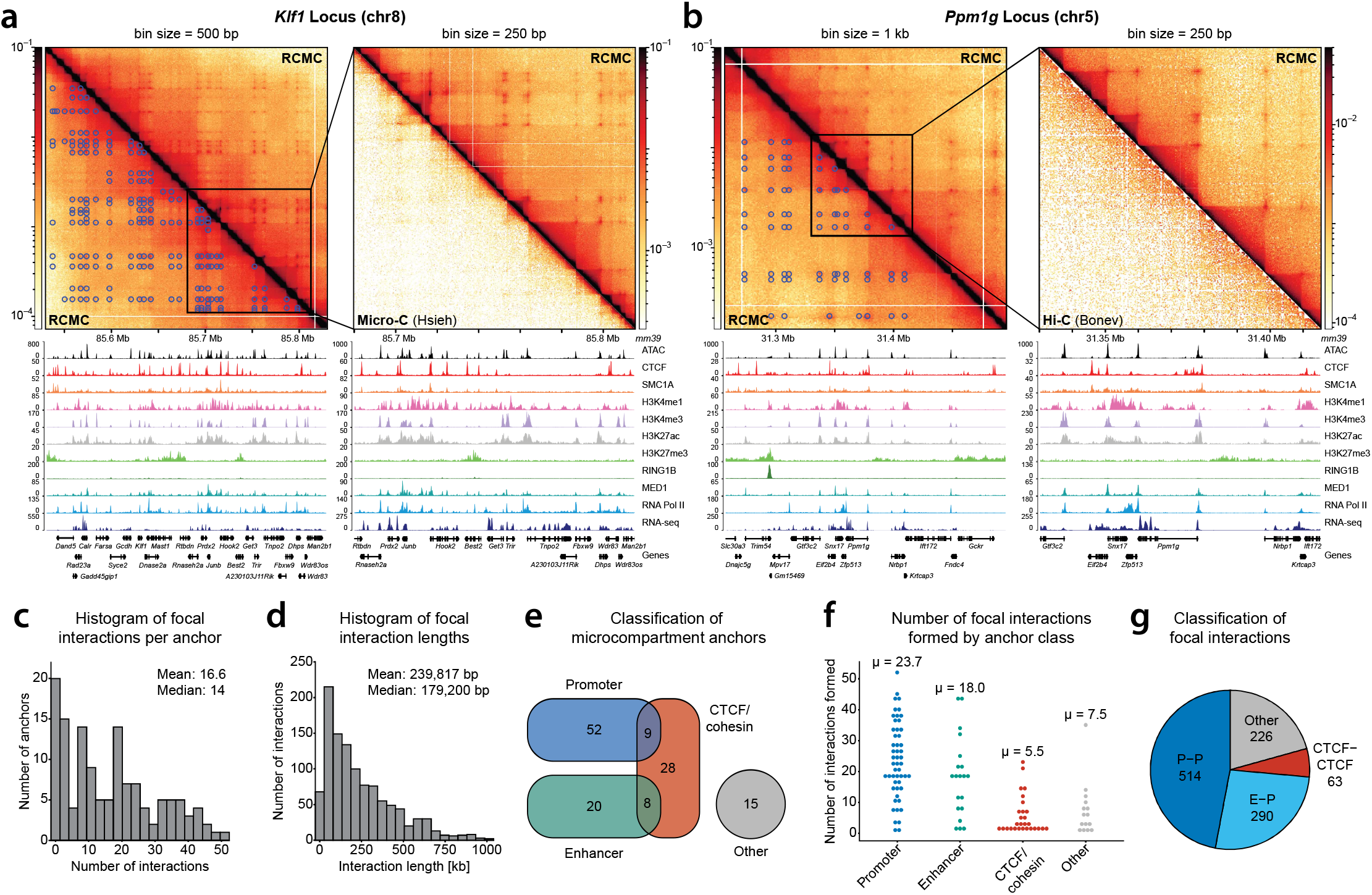
RCMC identifies highly nested, focal interactions called microcompartments which frequently connect enhancers and promoters. (**a-b**) Contact map visualization of RCMC data and called microcompartments at the *Klf1* (**a**) and *Ppm1g* (**b**) locus at 500 bp (**a**) and 1 kb (**b**) resolution (left) and 250 bp resolution (zoom in, right). Manually annotated microcompartment contacts are shown below the contact map diagonal on the left, while comparisons against genome-wide Micro-C^13^ (**a**) and Hi-C^31^ (**b**) are shown on the right. (**c-d**) Histograms showing distributions of (**c**) the number of focal interactions formed by microcompartment anchors and (**d**) the lengths spanned by focal interactions in kb. (**e**) Venn diagram of microcompartment anchor categories according to chromatin features overlapped by the enhancer ±1 kb. Promoters were defined as a region around annotated transcription start sites^50^ ±2 kb, enhancers as regions with overlapping peaks of in H3K4me1 (ENCFF282RLA) and H3K27ac (GSE90893) in ChIP-seq data which did not overlap promoters, and CTCF/cohesin as regions with overlapping peaks of CTCF (GSE90994) and SMC1A (GSE123636) in ChIP-seq data. Other regions are those not overlapping any of these features. (**f**) Swarm plot of the number of focal interactions formed by individual microcompartment anchors divided according to categories in (**e**), including the mean (µ) for each distribution. Anchors fitting into more than one category were excluded. (**g**) Fractions of loops classified into different categories: P-P (promoter-promoter), E-P (enhancer-promoter), CTCF-CTCF (CTCF/cohesin-CTCF/cohesin), other (Other-Other interactions, or any other combinations). CTCF-CTCF interactions do not include any anchors which overlap promoter or enhancer regions.

Next, we observed that these interactions resemble both loops and compartments. Like loops, they give rise to focal enrichments (“dots” in **Fig. 3a-b**) between two anchors and occasionally form contact domains as small as a few kilobases (“squares” in **Fig. 3a-b**). Like A/B-compartments, they result in nested interactions in a checkerboard-like fashion, with a mean of ∼17 interactions per anchor (mean loop length: ∼240 kb), and the most nested anchor forming more than 50 focal interactions (**Fig. 3c-d**). Because these highly nested and focal interactions (“dots”) resemble fine-scale compartmental interactions (see Discussion), we refer to them as *microcompartments*.

To understand which genomic elements form microcompartments, we investigated the chromatin states of microcompartment anchors (**Fig. 3c**; **Fig. S7**). About two-thirds of the identified microcompartment anchors overlapped either promoter (∼46%) or enhancer (∼21%) features (**Fig. 3e, Fig. S7**), with the remaining anchors either corresponding to CTCF and cohesin-bound anchors or unknowns (“Other”). Notably, however, promoters and enhancers formed many more focal interactions (**Fig. 3f**). Specifically, promoters and enhancers formed a mean of 24 and 18 interactions, respectively, compared to just 5.5 and 7.5 for CTCF and cohesin, and “other” anchors, respectively (**Fig. 3f**). Indeed, 74% of all annotated microcompartments represented either P-P or E-P interactions, while only 5% of interactions were between anchors which exclusively overlapped CTCF and cohesin (**Fig. 3g**). Taken together, these observations suggest that microcompartments largely represent nested interactions between promoter and enhancer regions as well as some currently poorly understood “other” regions.

### Microcompartment interactions are largely independent of cohesin and transcription

Having identified microcompartments as nested interactions frequently linking enhancers and promoters (**Fig. 3a-b**), we next took advantage of the cost-effective nature of RCMC to test the roles of loop extrusion and transcription in forming these interactions.

First, we explored the role of cohesin and cohesin-mediated loop extrusion. Acute loss of cohesin strengthens large-scale A/B compartments while at the same time causing the global loss of TADs, loop domains, and CTCF and cohesin-mediated structural loops^12,21,24,25,27,39^. Therefore, to understand whether cohesin is required for or antagonizes microcompartments, we used our previously validated mESC cell line to acutely deplete cohesin subunit RAD21 (mESC clone F1M RAD21-mAID-BFP-V5)^12,39^ and performed RCMC across all 5 regions with and without 3 hours of cohesin depletion (**Fig. 4a; Fig. S8a**). Cohesin depletion diminished the well-characterized CTCF and cohesin-mediated *Fbn2* loop^39^ (**Fig. S8a**), led to the expected change in contact frequency^21,23,24^ (**Fig. S8b**), and was reproducible between technical replicates (**Fig. S8c**), thus validating the cohesin depletion. As expected, the small fraction of interactions between CTCF and cohesin-bound sites showed large reductions in strength upon cohesin depletion (**Fig. 4a,c; Fig. S8a**). However, the strengths of other interaction types were largely unaffected by cohesin depletion. We therefore refine the microcompartment definition to interactions largely robust to cohesin depletion (see Discussion for full definition).

**Figure 4.**
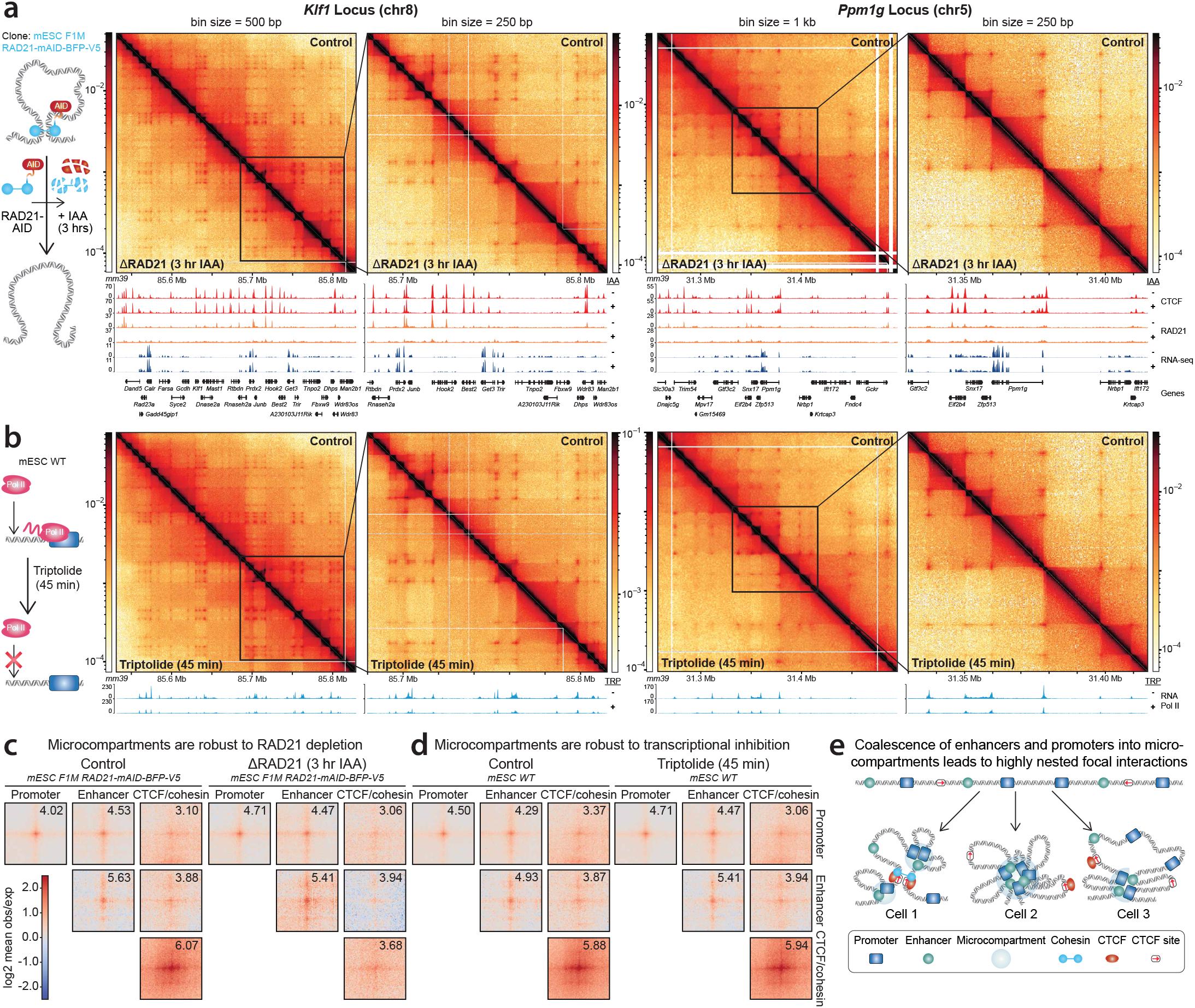
Microcompartments are largely robust to loss of loop extrusion and inhibition of transcription. (**a**) Cohesin (RAD21) depletion does not generally perturb microcompartments. Left: treatment paradigm for rapid depletion of RAD21 upon IAA treatment in clone F1M RAD21-mAID-BFP-V5 mESCs^12,39^. Right: Contact maps comparing a DMSO-treated control (above) and RAD21-depleted samples (below) are shown for the *Klf1* locus at 500 and 250 bp resolution and for the *Ppm1g* locus at 1 kb and 250 bp resolution. (**b**) Inhibition of transcription initiation with triptolide does not generally perturb microcompartments. Left: treatment paradigm for inhibition of transcription upon triptolide treatment in WT mESCs. Right: Contact maps comparing WT control (above) and transcriptionally-inhibited samples (below) are shown for the *Klf1* locus at 500 and 250 bp resolution and for the *Ppm1g* locus at 1 kb and 250 bp resolution. (**c-d**) Aggregate peak analysis matrix of called microcompartmental contacts showing loops across (**c**) RAD21 depletion and (**d**) transcriptional inhibition compared to their respective controls, separated by the identity of each contact’s constituent anchors. Plots show a 20 kb window centered on the loop at 250 bp resolution. Background-normalized intensity for a 1250×1250 bp box around the central dot for each aggregate peak is shown in the upper right of each plot. (**e**) Proposed model for the formation of microcompartments. Coalescence of multiple promoters and enhancer elements in a gene-dense region may occur through microphase separation as expected for an A/B block copolymer^4,44,45^, with strong A-A interactions. A/B block copolymer microphase separation is independent of loop extrusion. Block copolymer microphase separation may result in multiway interactions in different combinations of interactions being present in different cells, giving rise to the highly nested, focal interactions observed by RCMC, which averages across a population.

Second, we explored the role of transcription. We observed that microcompartments largely formed between active promoter and enhancer regions (**Fig. 3e; Fig. S7**), suggesting a relationship between active transcription and microcompartments. To understand if microcompartments are a downstream consequence of transcription, we abolished transcription by inhibiting transcription initiation by RNA Pol II using triptolide for 45 min^13,40^ and performed RCMC across all 5 regions (**Fig. 4b; Fig. S9**). Regardless of the type of interaction, microcompartment interactions were essentially unaffected by loss of transcription (**Fig. 4b, d**). We conclude that microcompartments do not require short-term transcription and are more likely either independent from or formed upstream of transcription rather than forming as a downstream consequence of transcription.

## DISCUSSION

Here we introduce RCMC as a new, accessible, and affordable method for mapping 3D genome structure at unprecedented depth. Compared with Micro-Capture-C^16^ methods such as TMCC^17^, RCMC is much more efficient (**Fig. 1b; Fig. S2c**), thus affording much higher depth with less sequencing. Another approach is to brute-force genome-wide Hi-C or Micro-C; by performing 150 separate Hi-C experiments and sequencing deeper than ever before, a recent preprint by Gu *et al*. reached 33 billion contacts^14^. However, such efforts^14^ are not accessible to most labs and poorly compatible with perturbation experiments vital to uncovering mechanisms of organization. Instead, with RCMC we reach the local equivalent of 200 billion contacts with relatively modest sequencing (**Fig. S2c**). We therefore propose RCMC as an ideal method for generating ultra-deep 3D contact maps and for perturbation experiments, albeit only for individual regions.

What molecular processes might drive microcompartment formation? Although cohesin-mediated loop extrusion is well-established to generate focal interactions (loops)^9,10^, microcompartmental loops are robust to acute cohesin removal, and therefore likely not dependent on loop extrusion (**Fig. 4a,c**). Furthermore, although most microcompartmental loops connect enhancers and promoters, microcompartments are also robust to acute to loss of RNA Pol II transcription initiation (**Fig. 4b,d**). Instead, we propose that nested and multiway focal microcompartments correspond to small A-compartments^14,41,42^ that form through a compartmentalization mechanism, perhaps mediated by factors upstream of RNA Pol II initiation, such as transcription factors and co-factors or active chromatin states^43^. Indeed, in the field of polymer physics, it is well-known that block copolymers undergo microphase separation^4,44,45^ when composed of distinct monomers that preferentially interact (**Fig. 4e**). Intuitively, if active chromatin regions at microcompartment anchors are selectively “sticky” with each other, they will tend to co-segregate, resulting in the formation of nested, focal interactions (**Fig. 4e**). Microphase separation due to preferential interactions among active loci within a block copolymer might thus explain the formation of the striking pattern of interactions we observe (**Fig. 3a-b; 4e**). In summary, we tentatively define microcompartments as follows: 1) highly nested, focal interactions that frequently connect promoters and enhancer regions often in gene-rich loci; 2) are formed through a compartmentalization mechanism; and 3) are largely independent of loop extrusion and transcription, at least on short timescales.

How do microcompartments compare to previously described 3D genome features? First, previous genome-wide Micro-C uncovered widespread short-range Promoter-Promoter and Enhancer-Promoter links (P-P and E-P links)^12,13^. Similarly, many microcompartment interactions connect promoters and enhancers. RCMC now better resolves these interactions, revealing them to be highly nested, frequently forming dozens of microcompartmental loops. Second, while differences in cell type preclude a direct comparison, the microcompartments described here also share features with the fine-scale A-compartment interactions recently described by Gu *et al*. that were proposed to segregate active enhancers and promoters into small A-compartments^14^. Along the lines of Gu *et al*., the microcompartments we observe form small contact domains, and their loops are more punctate as compared to CTCF and cohesin-mediated loops, which are more diffuse^14^ (**Fig. 4c-d**).

Finally, our study provides insights into the molecular mechanisms that mediate E-P interactions. While some studies proposed that cohesin is largely required for E-P interactions^27,46^, others have suggested that cohesin is most important for very long-range^47,48^ or for inducible E-P interactions^48,49^, or that cohesin is largely not required for the maintenance of E-P interactions^12,17^. Except for most CTCF and cohesin-bound enhancers and promoters, our data suggest that most P-P and E-P interactions are mediated by a compartmentalization mechanism distinct from loop extrusion. This may offer a mechanistic explanation for the observation that cohesin is not required for the short-term maintenance of most E-P interactions and that the effects of cohesin depletion on global gene expression are modest^12,17,25^.

In summary, we have introduced RCMC, uncovered microcompartments in gene-rich areas in mESCs, and shown that microcompartments require neither cohesin nor transcription. RCMC provides an accessible method to deeply resolve 3D genome structure in general and enhancer-promoter interactions in particular across loci, cell types, and disease states. In the future, it will be important to test the generality of microcompartments, to further dissect their molecular basis and regulation, and to understand the frequency and lifetime of microcompartmental interactions in live cells^39^.

## Acknowledgements

We thank TH Stanley Hsieh, the co-inventor of Micro-C, for holding a Micro-C workshop to teach us the protocol. We also thank Stanley, Claudia Cattoglio, and Leonid Mirny for extensive and insightful discussions throughout this project. We thank Leonid Mirny and Geoff Fudenberg for insightful discussions on ICE normalization. We thank Job Dekker, Simon Grosse-Holz, Emily Navarette, Sameer Abraham, and the Hansen lab for helpful discussions throughout this project. We thank Elphege Nora, Miriam Huntley, Alistair Boettiger, Stanley Hsieh, Domenic Narducci, Sarah Nemsick, Asmita Jha, Michele Gabriele, Jin Harvey Yang, Christine Tyler, Sarah Johnstone, Christian Cerda-Smith, and Vijay Sankaran for critical feedback on the manuscript. We thank Marieke Oudelaar for providing TMCC numbers and feedback on the TMCC vs. RCMC comparison and the manuscript. We thank the Koch Institute’s Robert A. Swanson (1969) Biotechnology Center for technical support, specifically the Integrated Genomics and Bioinformatics Core and MIT BioMicroCenter, and this work was supported in part by the Koch Institute Support (core) Grant P30-CA14051 from the National Cancer Institute. We also thank the Walk-Up Sequencing services of the Broad Institute of MIT and Harvard. This work was supported by NIH grants DP2GM140938, R33CA257878, and UM1HG011536, NSF grant 2036037, a Solomon Buchsbaum Research Support Committee award and the Koch Institute Frontier Research Fund. MKH is supported by an NIH F32GM140548 fellowship and a non-stipendiary EMBO fellowship. The raw data can be found at NCBI GEO under accession number GSE207225.

## REFERENCES

1. Dekker, J. et al. The 3D Genome as Moderator of Chromosomal Communication. Cell (2016). doi:10.1016/j.cell.2016.02.007

2. Oudelaar, A. M. et al. The relationship between genome structure and function. Nat. Rev. Genet. 22, 154–168 (2021).

3. Lieberman-Aiden, E. et al. Comprehensive Mapping of Long-Range Interactions Reveals Folding Principles of the Human Genome. Science (80-.). 326, 289 LP – 293 (2009).

4. Nuebler, J. et al. Chromatin organization by an interplay of loop extrusion and compartmental segregation. Proc. Natl. Acad. Sci. 115, E6697 LP–E6706 (2018).

5. Dixon, J. R. et al. Topological domains in mammalian genomes identified by analysis of chromatin interactions. Nature 485, 376– 380 (2012).

6. Nora, E. P. et al. Spatial partitioning of the regulatory landscape of the X-inactivation centre. Nature 485, 381–385 (2012).

7. Rao, S. S. P. et al. A 3D Map of the Human Genome at Kilobase Resolution Reveals Principles of Chromatin Looping. Cell 159, 1665–1680 (2014).

8. Goel, V. Y. et al. The macro and micro of chromosome conformation capture. WIREs Dev. Biol. n/a, e395 (2020).

9. Sanborn, A. L. et al. Chromatin extrusion explains key features of loop and domain formation in wild-type and engineered genomes. Proc. Natl. Acad. Sci. U. S. A. (2015). doi:10.1073/pnas.1518552112

10. Fudenberg, G. et al. Formation of Chromosomal Domains by Loop Extrusion. Cell Rep. (2016). doi:10.1016/j.celrep.2016.04.085

11. Krietenstein, N. et al. Ultrastructural Details of Mammalian Chromosome Architecture. Mol. Cell (2020). doi:https://doi.org/10.1016/j.molcel.2020.03.003

12. Hsieh, T.-H. S. et al. Enhancer-promoter interactions and transcription are maintained upon acute loss of CTCF, cohesin, WAPL, and YY1. bioRxiv 2021.07.14.452365 (2021). doi:10.1101/2021.07.14.452365

13. Hsieh, T.-H. S. et al. Resolving the 3D Landscape of Transcription-Linked Mammalian Chromatin Folding. Mol. Cell (2020). doi:https://doi.org/10.1016/j.molcel.2020.03.002

14. Gu, H. et al. Fine-mapping of nuclear compartments using ultra-deep Hi-C shows that active promoter and enhancer elements localize in the active A compartment even when adjacent sequences do not. bioRxiv 2021.10.03.462599 (2021). doi:10.1101/2021.10.03.462599

15. Hansen, A. S. et al. Distinct Classes of Chromatin Loops Revealed by Deletion of an RNA-Binding Region in CTCF. Mol. Cell (2019). doi:https://doi.org/10.1016/j.molcel.2019.07.039

16. Hua, P. et al. Defining genome architecture at base-pair resolution. Nature 595, 125–129 (2021).

17. Aljahani, A. et al. Analysis of sub-kilobase chromatin topology reveals nano-scale regulatory interactions with variable dependence on cohesin and CTCF. Nat. Commun. 13, 2139 (2022).

18. Gasperini, M. et al. A Genome-wide Framework for Mapping Gene Regulation via Cellular Genetic Screens. Cell 176, 377-390.e19 (2019).

19. Barshad, G. et al. RNA polymerase II and PARP1 shape enhancer-promoter contacts. bioRxiv 2022.07.07.499190 (2022). doi:10.1101/2022.07.07.499190

20. Zhang, S. et al. Enhancer-promoter contact formation requires RNAPII and antagonizes loop extrusion. bioRxiv 2022.07.04.498738 (2022). doi:10.1101/2022.07.04.498738

21. Schwarzer, W. et al. Two independent modes of chromatin organization revealed by cohesin removal. Nature (2017). doi:10.1038/nature24281

22. Nora, E. P. et al. Targeted Degradation of CTCF Decouples Local Insulation of Chromosome Domains from Genomic Compartmentalization. Cell (2017). doi:10.1016/j.cell.2017.05.004

23. Gassler, J. et al. A mechanism of cohesin-dependent loop extrusion organizes zygotic genome architecture. EMBO J. 36, 3600–3618 (2017).

24. Wutz, G. et al. Topologically associating domains and chromatin loops depend on cohesin and are regulated by CTCF, WAPL, and PDS5 proteins. EMBO J. (2017). doi:10.15252/embj.201798004

25. Rao, S. S. P. et al. Cohesin Loss Eliminates All Loop Domains. Cell (2017). doi:10.1016/j.cell.2017.09.026

26. Haarhuis, J. H. I. et al. The Cohesin Release Factor WAPL Restricts Chromatin Loop Extension. Cell 169, 693-707.e14 (2017).

27. El Khattabi, L. et al. A Pliable Mediator Acts as a Functional Rather Than an Architectural Bridge between Promoters and Enhancers. Cell 178, 1145-1158.e20 (2019).

28. Oudelaar, A. M. et al. Dynamics of the 4D genome during in vivo lineage specification and differentiation. Nat. Commun. 11, 2722 (2020).

29. Jäger, R. et al. Capture Hi-C identifies the chromatin interactome of colorectal cancer risk loci. Nat. Commun. 6, 6178 (2015).

30. Imakaev, M. et al. Iterative correction of Hi-C data reveals hallmarks of chromosome organization. Nat. Methods 9, 999–1003 (2012).

31. Bonev, B. et al. Multiscale 3D Genome Rewiring during Mouse Neural Development. Cell 171, 557-572.e24 (2017).

32. Zhou, H. Y. et al. A Sox2 distal enhancer cluster regulates embryonic stem cell differentiation potential. Genes Dev. 28, 2699– 2711 (2014).

33. Li, Y. et al. CRISPR Reveals a Distal Super-Enhancer Required for Sox2 Expression in Mouse Embryonic Stem Cells. PLoS One 9, e114485 (2014).

34. Chakraborty, S. et al. High affinity enhancer-promoter interactions can bypass CTCF/cohesin-mediated insulation and contribute to phenotypic robustness. bioRxiv 2021.12.30.474562 (2022). doi:10.1101/2021.12.30.474562

35. Akgol Oksuz, B. et al. Systematic evaluation of chromosome conformation capture assays. Nat. Methods 18, 1046–1055 (2021).

36. Roayaei Ardakany, A. et al. Mustache: multi-scale detection of chromatin loops from Hi-C and Micro-C maps using scale-space representation. Genome Biol. 21, 256 (2020).

37. Sergey Venev, Nezar Abdennur, Anton Goloborodko, Ilya Flyamer, Geoffrey Fudenberg, Johannes Nuebler, Aleksandra Galitsyna, Betul Akgol, Sameer Abraham, Peter Kerpedjiev, & M. I. open2c/cooltools: v0.4.1 (v0.4.1). Zenodo https://doi.org/10.5281/zenodo.5214125 (2021). xdoi:https://doi.org/10.5281/zenodo.5214125

38. Abdennur, N. et al. Cooler: scalable storage for Hi-C data and other genomically labeled arrays. Bioinformatics 36, 311–316 (2020).

39. Gabriele, M. et al. Dynamics of CTCF- and cohesin-mediated chromatin looping revealed by live-cell imaging. Science (80-.). 376, 496–501 (2022).

40. Jonkers, I. et al. Genome-wide dynamics of Pol II elongation and its interplay with promoter proximal pausing, chromatin, and exons. Elife 3, e02407 (2014).

41. Rosencrance, C. D. et al. Chromatin Hyperacetylation Impacts Chromosome Folding by Forming a Nuclear Subcompartment. Mol. Cell 78, 112-126.e12 (2020).

42. You, Q. et al. Direct DNA crosslinking with CAP-C uncovers transcription-dependent chromatin organization at high resolution. Nat. Biotechnol. 39, 225–235 (2021).

43. Rippe, K. et al. Functional organization of RNA polymerase II in nuclear subcompartments. Curr. Opin. Cell Biol. 74, 88–96 (2022).

44. Leibler, L. Theory of Microphase Separation in Block Copolymers. Macromolecules 13, 1602–1617 (1980).

45. Meier, D. J. Theory of block copolymers. I. Domain formation in A-B block copolymers. J. Polym. Sci. Part C Polym. Symp. 26, 81–98 (1969).

46. Thiecke, M. J. et al. Cohesin-Dependent and -Independent Mechanisms Mediate Chromosomal Contacts between Promoters and Enhancers. Cell Rep. 32, (2020).

47. Kane, L. et al. Cohesin is required for long-range enhancer action. bioRxiv 2021.06.24.449812 (2021). doi:10.1101/2021.06.24.449812

48. Calderon, L. et al. Cohesin-dependence of neuronal gene expression relates to chromatin loop length. Elife 11, e76539 (2022).

49. Cuartero, S. et al. Control of inducible gene expression links cohesin to hematopoietic progenitor self-renewal and differentiation. Nat. Immunol. 19, 932–941 (2018).

50. Navarro Gonzalez, J. et al. The UCSC Genome Browser database: 2021 update. Nucleic Acids Res. 49, D1046–D1057 (2021).

