## Supplementary Materials for "Region Capture Micro-C reveals coalescence of enhancers and promoters into nested microcompartments"

#### Contents

- Supplementary Figures and Figure Legends
- Supplementary Table
- Materials and Methods
- Supplementary References
- Region Capture Micro-C protocol

### Supplementary Figures and Figure Legends

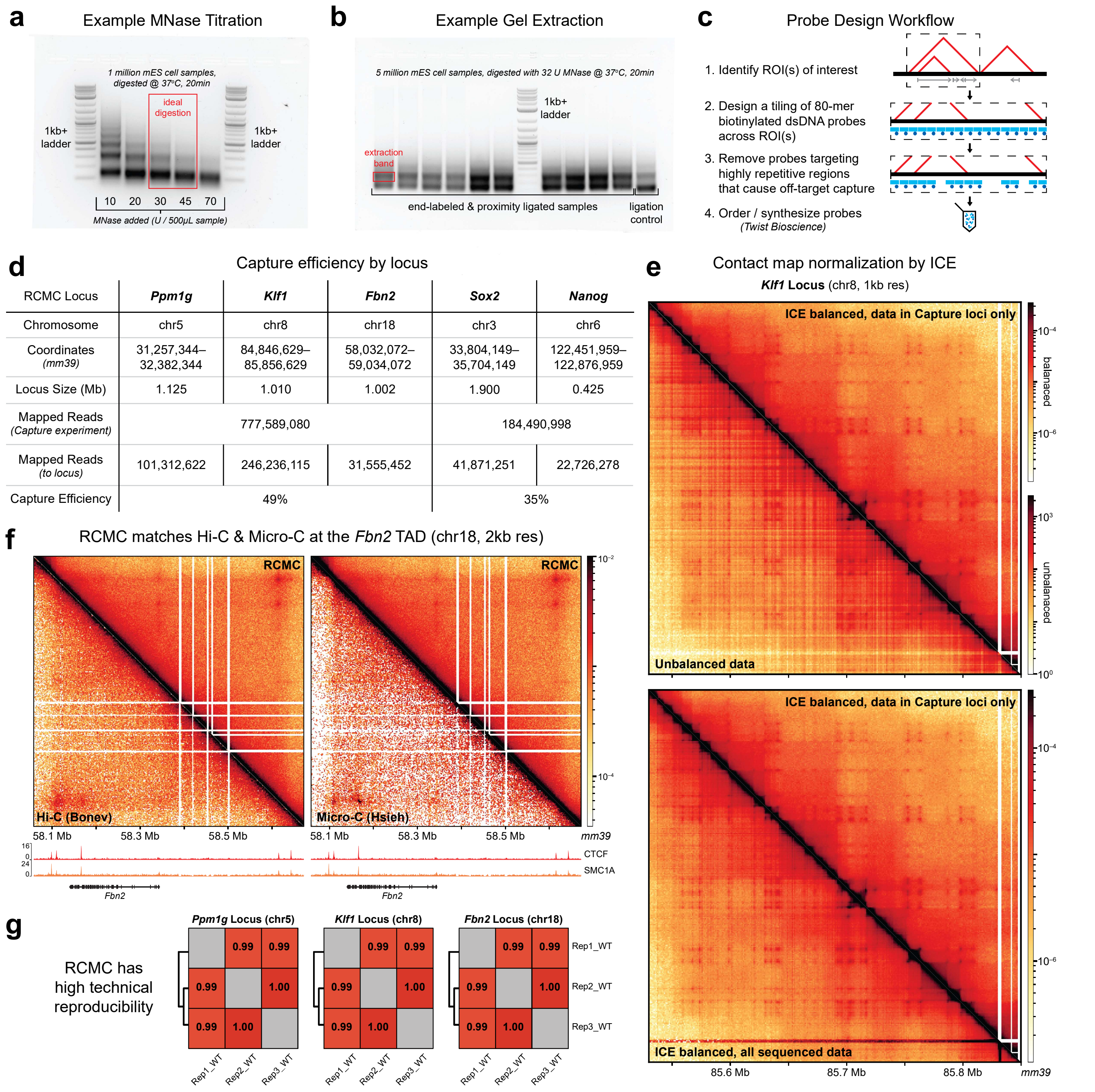


**Supplementary Figure 1. RCMC efficiently and reproducibly captures ligated dinucleosomal fragments, giving rise to deep contact maps.** (**a**) Representative MNase titration DNA gel indicating the ideal level of digestion by MNase, based on the ratio of mononucleosomal/dinucleosomal fragments, for the RCMC protocol. (**b**) Representative size selection gel for the RCMC protocol showing the necessary separation and the dinucleosomal band that is extracted to obtain ligated fragments. (**c**) Overview of the capture probe design workflow for RCMC. 80mer probes tiling the region of interest are designed, removing those which overlap highly repetitive regions. (**d**) Summary of the capture efficiency for each of the five regions for which probes were designed. The locations and sizes of the regions, the number of ligated fragments which mapped at single loci at both ends in total and in the region, and the capture efficiencies are given. (**e**) Contact maps comparing raw, unbalanced data (upper panel, lower triangle), ICE[1] balanced to all aligned reads (lower panel, lower triangle) and ICE balanced to reads in Capture loci only (both panels, upper triangle). Balancing only to data within the Capture loci was necessary to achieve a contact map without artefacts due to capture bias. (**f**) Contact maps comparing the entire *Fbn2* TAD in RCMC and in Hi-C[2] and Micro-C[3]. Gene annotations and ChIP-seq signal tracks are shown below the contact maps. (**g**) Measurement of reproducibility between technical replicates at the *Ppm1g*, *Klf1*, and *Fbn2* loci, with reproducibility scores determined using HiCRep[4], clustered according to similarity.

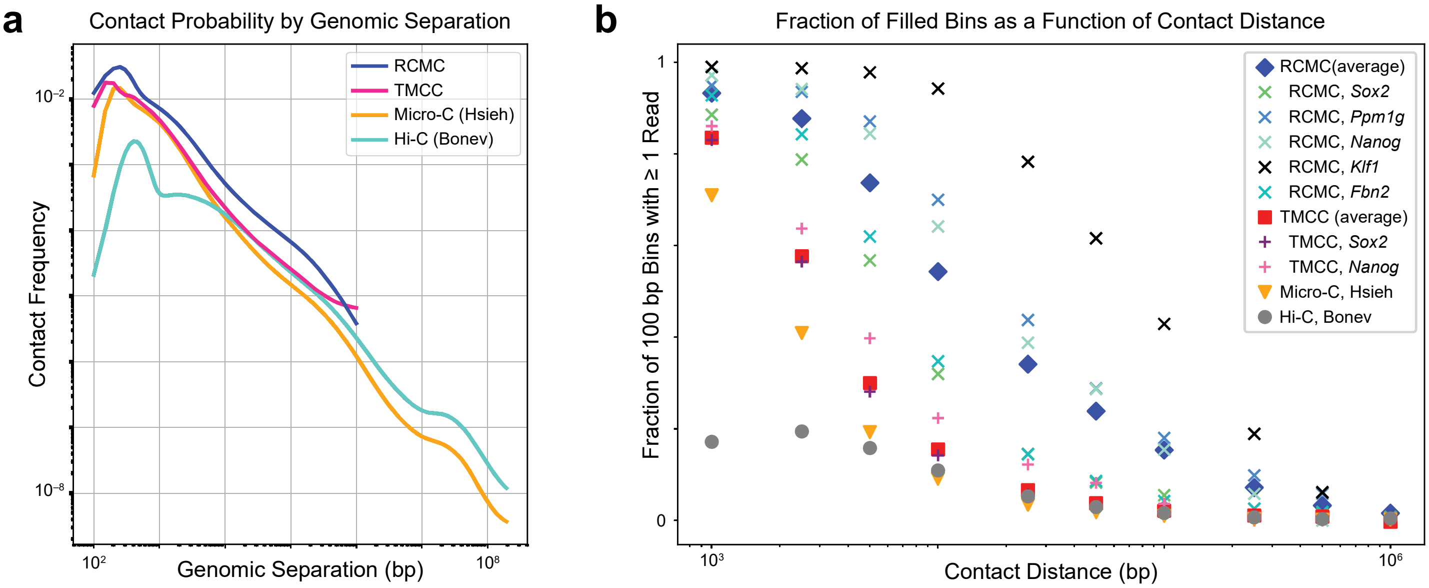


**c** Comparison of Structurally Informative Contacts Across Relevant 3C Methods

| Read Counts Comparison | Locus size (Mb) | Mapped sequencing reads (total) | Mapped sequencing reads (unique) Read count | Mapped sequencing reads (unique) % total | Unique contacts, >= 1 kb Read count | Unique contacts, >= 1 kb % unique |
|---|---|---|---|---|---|---|
| Region Capture Micro-C (RCMC) | N/A | 962,080,078 | 542,659,346 | 56 | 433,866,124 | 80 |
| *RCMC, Sox2 locus (chr3)* | 1.900 | 41,871,251 | 18,932,557 | 45 | 16,231,438 | 86 |
| *RCMC, Ppm1g locus (chr5)* | 1.125 | 101,312,622 | 37,856,541 | 37 | 32,430,599 | 86 |
| *RCMC, Nanog locus (chr6)* | 0.425 | 22,726,278 | 10,234,592 | 45 | 8,417,054 | 82 |
| *RCMC, Klf1 locus (chr8)* | 1.010 | 246,236,115 | 80,716,455 | 33 | 66,569,425 | 82 |
| *RCMC, Fbn2 locus (chr18)* | 1.002 | 31,555,452 | 12,802,056 | 41 | 11,142,511 | 87 |
| Tiled-Micro-Capture-C (TMCC), Aljahani et al. (2022) | N/A | 1,120,446,062 | 233,233,787 | 19 | 8,819,602<br>13,402,554* | 4<br>50* |
| *TMCC, Sox2 locus (chr3)* | 1.280 | 396,415,581 | 34,524,304 | 9 | 4,327,272<br>6,730,085* | 13<br>52* |
| *TMCC, Nanog locus (chr6)* | 0.250 | 73,941,806 | 6,295,110 | 9 | 1,384,329<br>1,740,491* | 22<br>50* |
| Micro-C, Hsieh et al. (2020) | N/A | 2,918,350,701 | 2,644,025,735 | 91 | 2,105,436,125 | 80 |
| Hi-C, Bonev et al. (2017) | N/A | 7,260,480,082 | 3,333,795,091 | 46 | 3,200,749,372 | 96 |

**Supplementary Figure 2. Benchmarking of RCMC against other 3C methods.** (**a**) Contact probability curves comparing RCMC against the highest resolution Tiled-Micro-Capture-C (TMCC)[5], Micro-C[3], and Hi-C[2] mESC datasets across contact distances. (**b**) Benchmarking comparison of RCMC's ability to fill out high-resolution contact matrices against TMCC[5], Micro-C[3], and Hi-C[2]. Region-averaged calculations are shown for RCMC, TMCC, Micro-C, and Hi-C, and calculations for individual captured regions are also shown for RCMC and TMCC. The x-axis shows the contact distance in bp, and the y-axis shows the fraction of all 100 bp contact bins at a given contact distance within the Captured locus that contain at least one read. (**c**) Summary of read counts across RCMC, TMCC[5], Micro-C[3], and Hi-C[2]. The total number of mapped sequencing reads, the fraction of those reads that are unique, and the fraction of unique reads that are structurally informative (defined as *cis* contacts >=1 kb) are given for each method. For the TMCC numbers, two versions are provided. In black, we show numbers processed by us using the same bioinformatic pipeline as for RCMC. Capture region-specific quantifications (defined here as all reads with at least one of two read mates mapped to the locus) are also provided for all RCMC loci and the *Sox2* and *Nanog* TMCC loci; the *Oct4* and *Prdm14* TMCC loci are not considered in this manuscript. In red are shown numbers kindly provided by Dr. A Marieke Oudelaar, obtained using the custom TMCC-specific bioinformatic pipeline from Aljahani *et al.*[5].Values in red with asterisks denote quantifications of all unique contact pairs mapped to Captured loci (not filtered to be >= 1 kb in size).

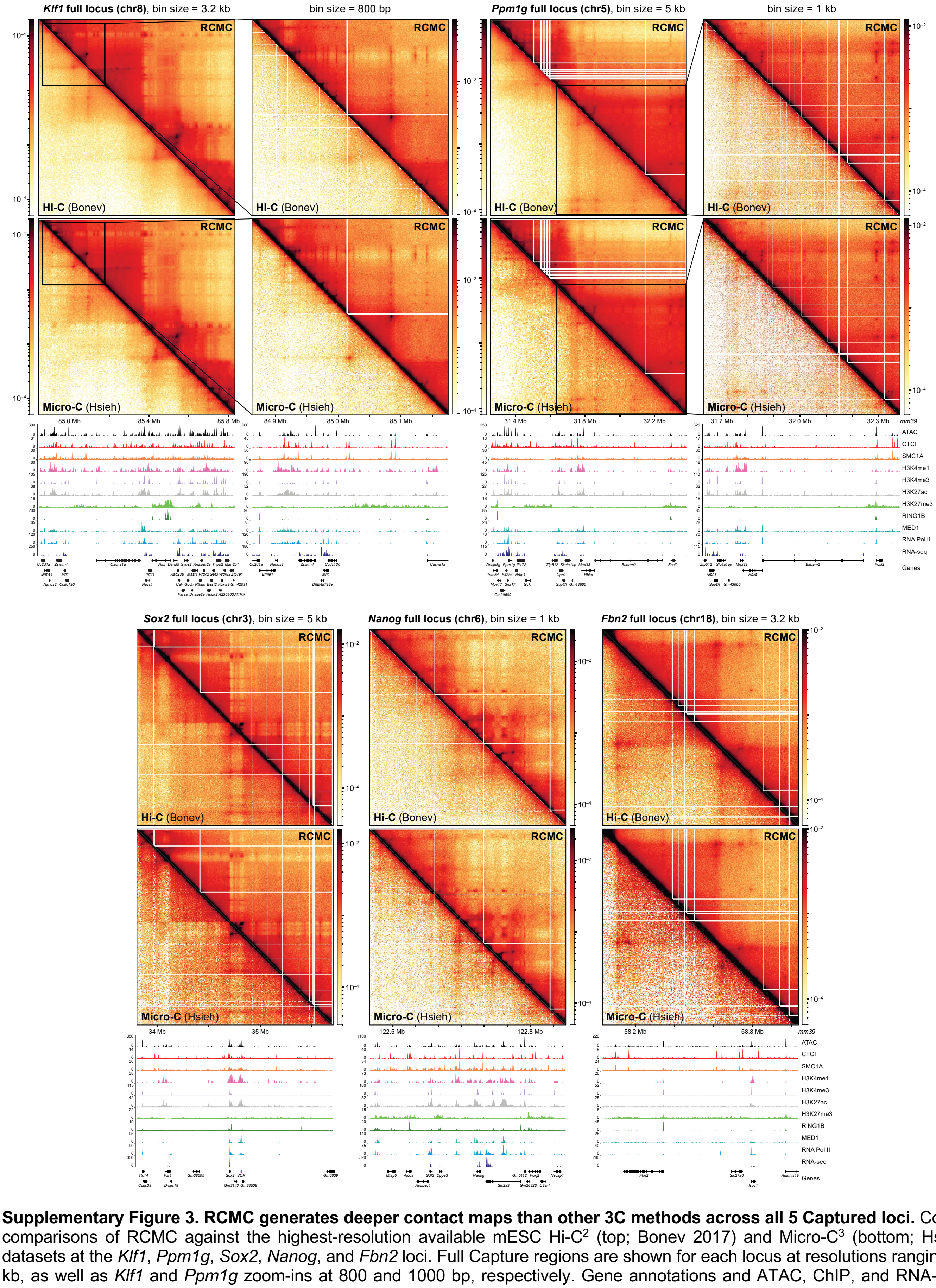


**Supplementary Figure 3. RCMC generates deeper contact maps than other 3C methods across all 5 Captured loci.** Contact map comparisons of RCMC against the highest-resolution available mESC Hi-C[2] (top; Bonev 2017) and Micro-C[3] (bottom; Hsieh 2020) datasets at the *Klf1*, *Ppm1g*, *Sox2*, *Nanog*, and *Fbn2* loci. Full Capture regions are shown for each locus at resolutions ranging from 1-5 kb, as well as *Klf1* and *Ppm1g* zoom-ins at 800 and 1000 bp, respectively. Gene annotations and ATAC, ChIP, and RNA-seq tracks (Supplementary Table 1) are shown below the contact maps, while the contact intensity scales are shown next to the maps.

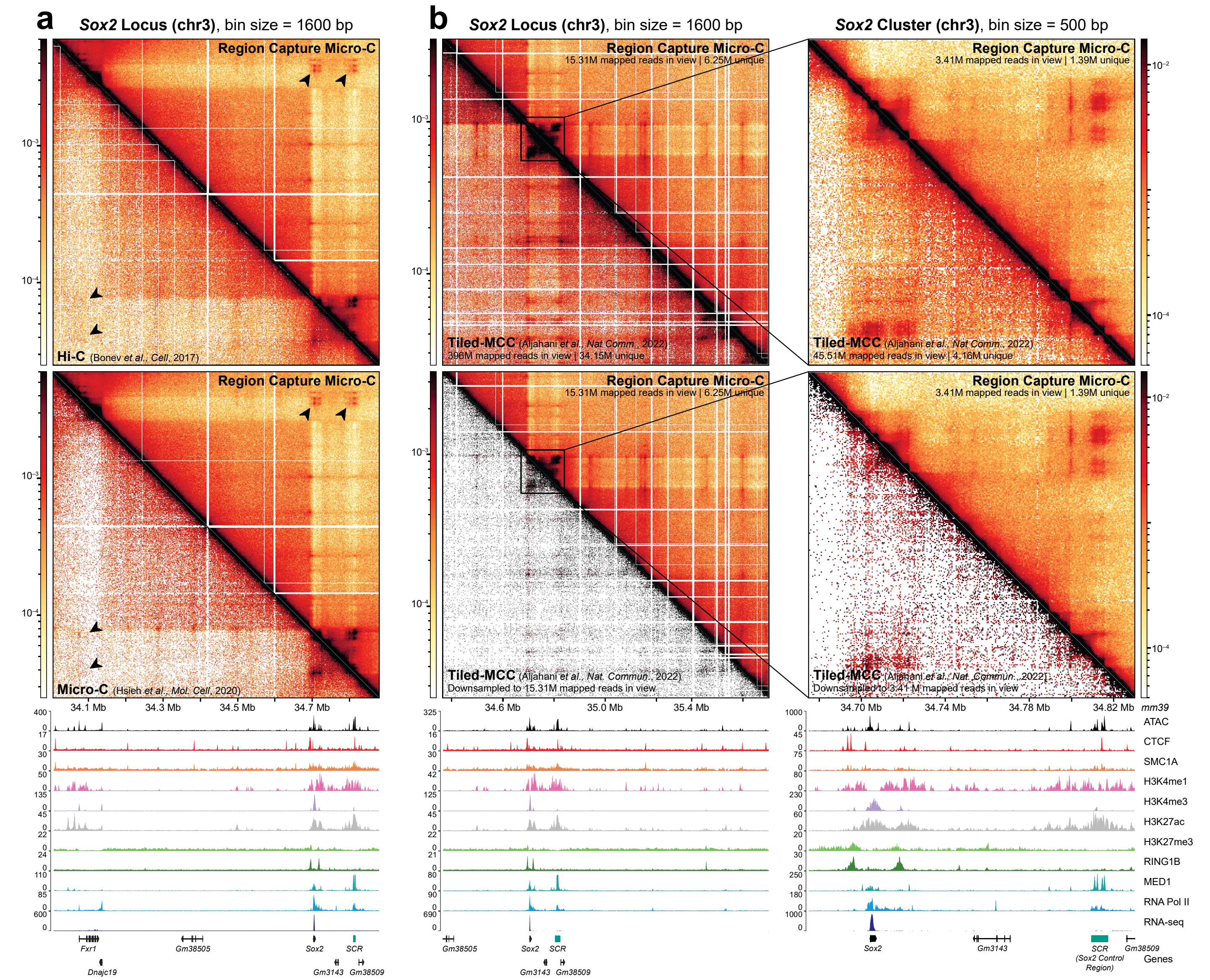


**Supplementary Figure 4. RCMC maps the *Sox2* locus more deeply and efficiently than sister methods, uncovering novel interactions.** (**a**) Contact map comparisons of RCMC against Hi-C[2] (top) and Micro-C[3] (bottom) at the *Sox2* locus at 1.6 kb resolution. Arrows mark contacts between *Sox2*, the SCR, and *Fxr1* not mapped by Hi-C and Micro-C. (**b**) Contact map comparisons of RCMC against Tiled-Micro-Capture-C[5] (TMCC) across the whole TMCC-Captured locus (left, 1600 bp resolution) and in the *Sox2* and SCR regulatory cluster (right, 500 bp resolution). Full datasets are visualized in the top contact maps, and TMCC has been downsampled to match the total number of RCMC sequencing reads in view in the bottom contact maps.

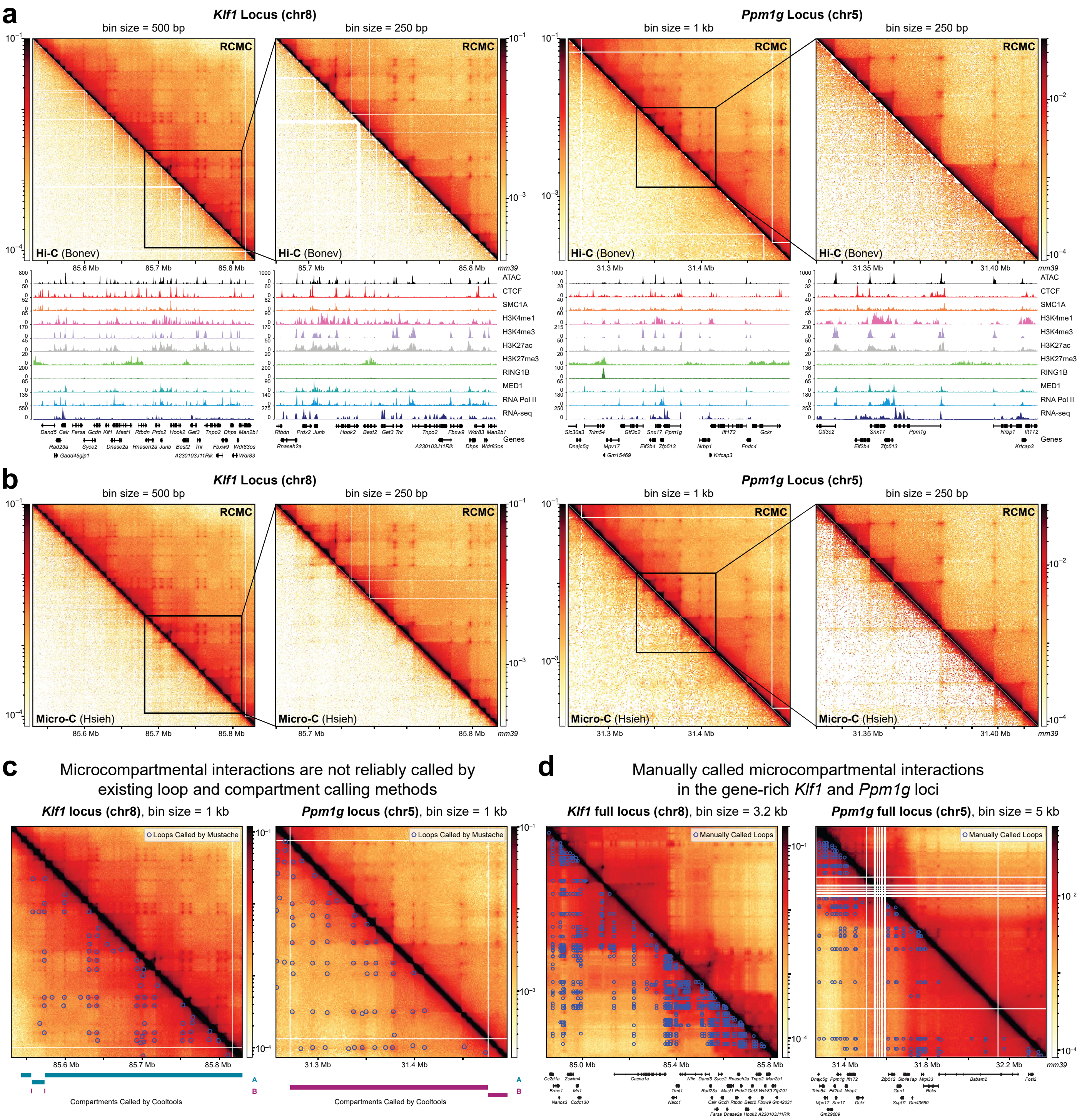


**Supplementary Figure 5. RCMC identifies microcompartments, which are not visible in other methods and not reliably called by existing algorithms.** (**a-b**) Contact maps comparison of RCMC (top) against Hi-C[2] (bottom, **a**) and Micro-C[3] (bottom, **b**) at the *Klf1* locus at 500 and 250 bp resolutions and at the *Ppm1g* locus at 1000 and 250 bp resolutions. (**c**) Contact map of the *Klf1* locus with loop calls by Mustache[6] at 1 kb resolution overlaid on the bottom half of the map and compartment calls by cooltools[7,8] shown below the map. (**d**) Contact maps of the entire *Klf1* (3.2 kb resolution) and *Ppm1g* (5 kb resolution) Captured loci with manually called loops (see Methods) overlaid on the bottom halves of the maps.

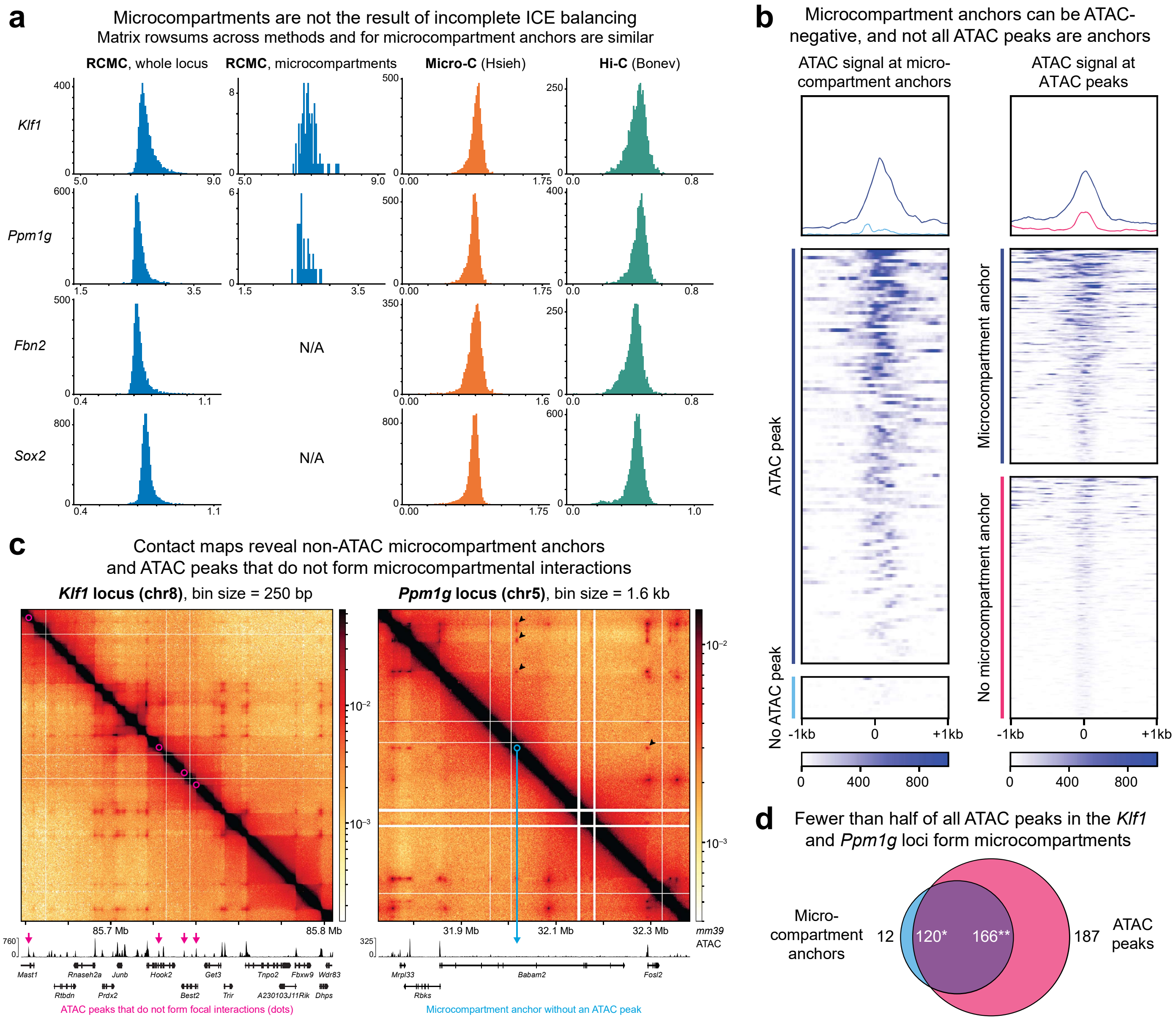


**Supplementary Figure 6. Microcompartments are not artifacts resulting from incomplete ICE balancing nor chromatin accessibility.** (**a**) Comparison of ICE balancing across methods and Capture loci. Distributions of the sums of ICE-balanced contact matrix rows at 250 bp resolution are shown at the *Klf1*, *Ppm1g*, *Fbn2*, and *Sox2* loci for RCMC, Micro-C[3], and Hi-C[2], as well as for the subset of RCMC rows containing microcompartment anchors. A sharp unimodal peak is consistent with ICE's baseline assumption that all contact matrix rows and columns must sum to the same value. (**b**) Metaplots (above) and heatmaps (below) depicting ATAC signal at microcompartment anchors (left, separated by whether anchors coincide with an ATAC peak) and at all ATAC peaks in the *Klf1* and *Ppm1g* Capture loci (right, separated by whether peaks coincide with a microcompartment anchor). Signals are plotted in a 2 kb window centered on the anchor (left) or the ATAC peak (right). (**c**) RCMC contact maps at the *Klf1* (left, 250 bp resolution) and *Ppm1g* (right, 1.6 kb resolution) indicating ATAC peaks that do not form microcompartments (left, magenta) and a microcompartment anchor that does not coincide with an ATAC peak (right, cyan). Black arrows (right) indicate microcompartmental loops involving the ATAC-negative microcompartment anchor. (**d**) Venn diagram breakdown of the overlap between all manually annotated microcompartment anchors and all ATAC peaks across the *Klf1* and *Ppm1g* Capture loci. Of 132 annotated microcompartment anchors, 12 do not coincide with ATAC peaks (cyan) while 120 do (purple, *). Of 353 called ATAC peaks, 187 do not form microcompartment anchors (magenta) while 166 do (purple, **). The apparent discrepancy of 120 microcompartment anchors being anchored by 166 ATAC peaks is due to two close ATAC peaks occasionally anchoring a single microcompartment.

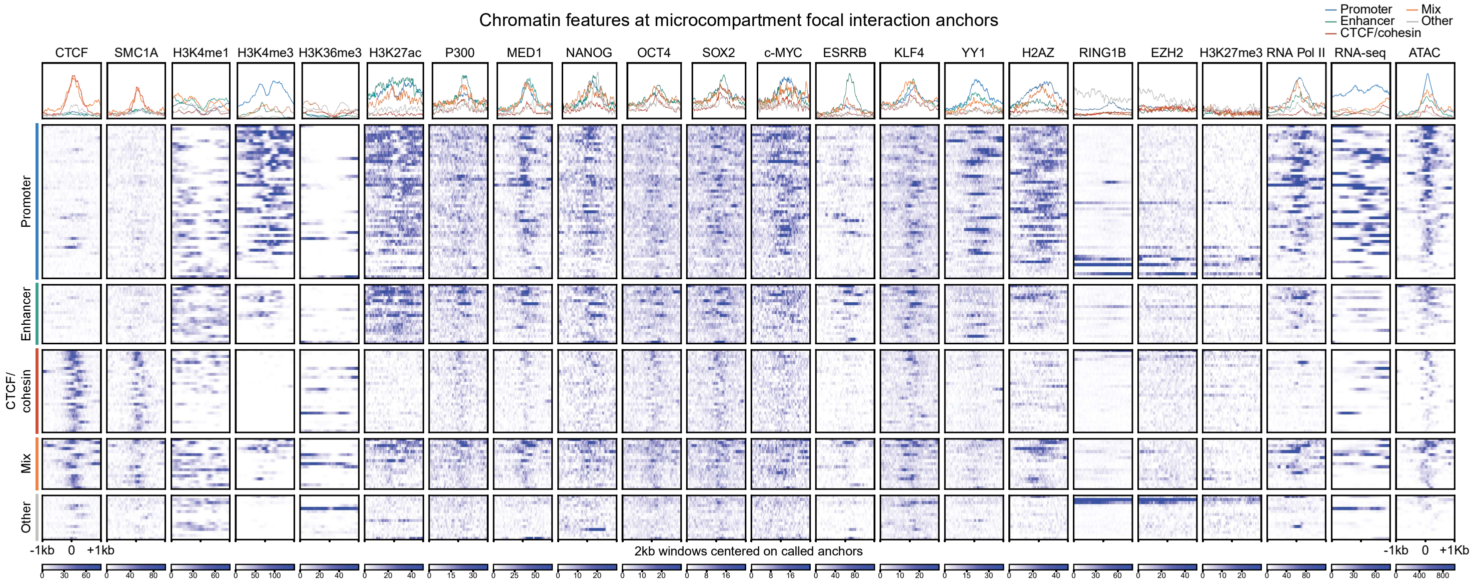


**Supplementary Figure 7. Categories of microcompartment anchors can be defined by their chromatin features.** Metaplots (above) and heatmaps (below) depicting ATAC, ChIP, and RNA-seq (Supplementary Table 1) signal at microcompartment loop anchors for classes of microcompartment anchors as defined in Fig. 3e. Features are plotted in a 2 kb window centered on the anchor.

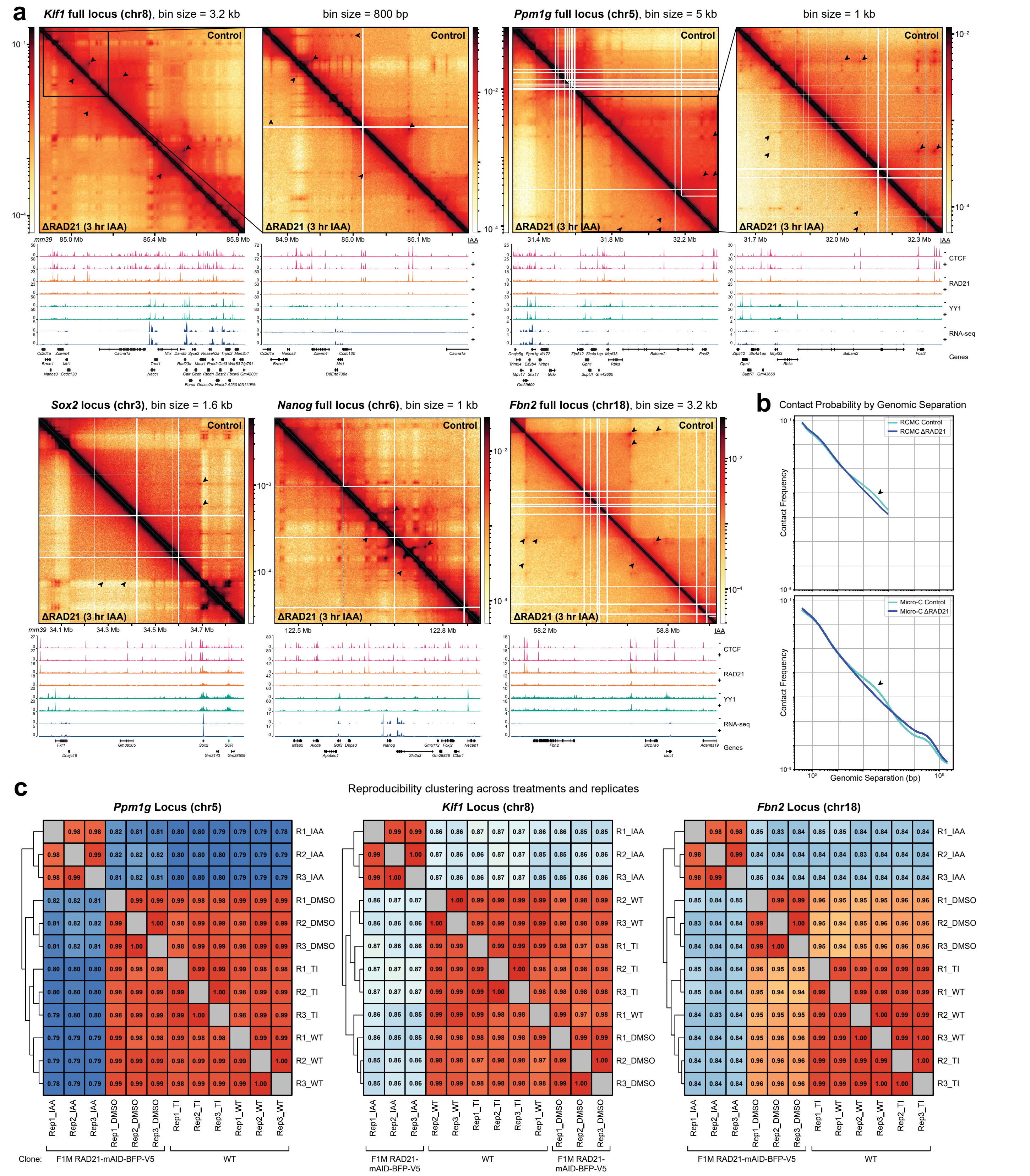


**Supplementary Figure 8. Cohesin depletion disrupts CTCF/Cohesin loops, but generally not microcompartmental loops.** (**a**) Contact maps comparing a DMSO control (above) and RAD21-depleted samples (below) are shown for the *Klf1*, *Ppm1g*, *Sox2*, *Nanog*, and *Fbn2* loci at resolutions spanning 800 bp – 5 kb in F1M RAD21-mAID-BFP-V5 mESCs[9,10]. Arrows mark contacts lost upon RAD21 depletion. ChIP data from Hsieh et al., *bioRxiv* (2021)[10] is shown below the maps before and after the IAA treatment (500 µM, 3 hours). (**b**) Contact probability curves comparing RAD21-depleted RCMC samples against a DMSO control (top) RAD21-depleted Micro-C samples against a DMSO control (bottom). Arrows indicate the contact frequency "bump" lost upon RAD21 depletion. (**c**) Measurement of reproducibility between technical replicates and across treatments at the *Ppm1g*, *Klf1*, and *Fbn2* loci, with reproducibility scores determined using HiCRep[4], clustered according to similarity.

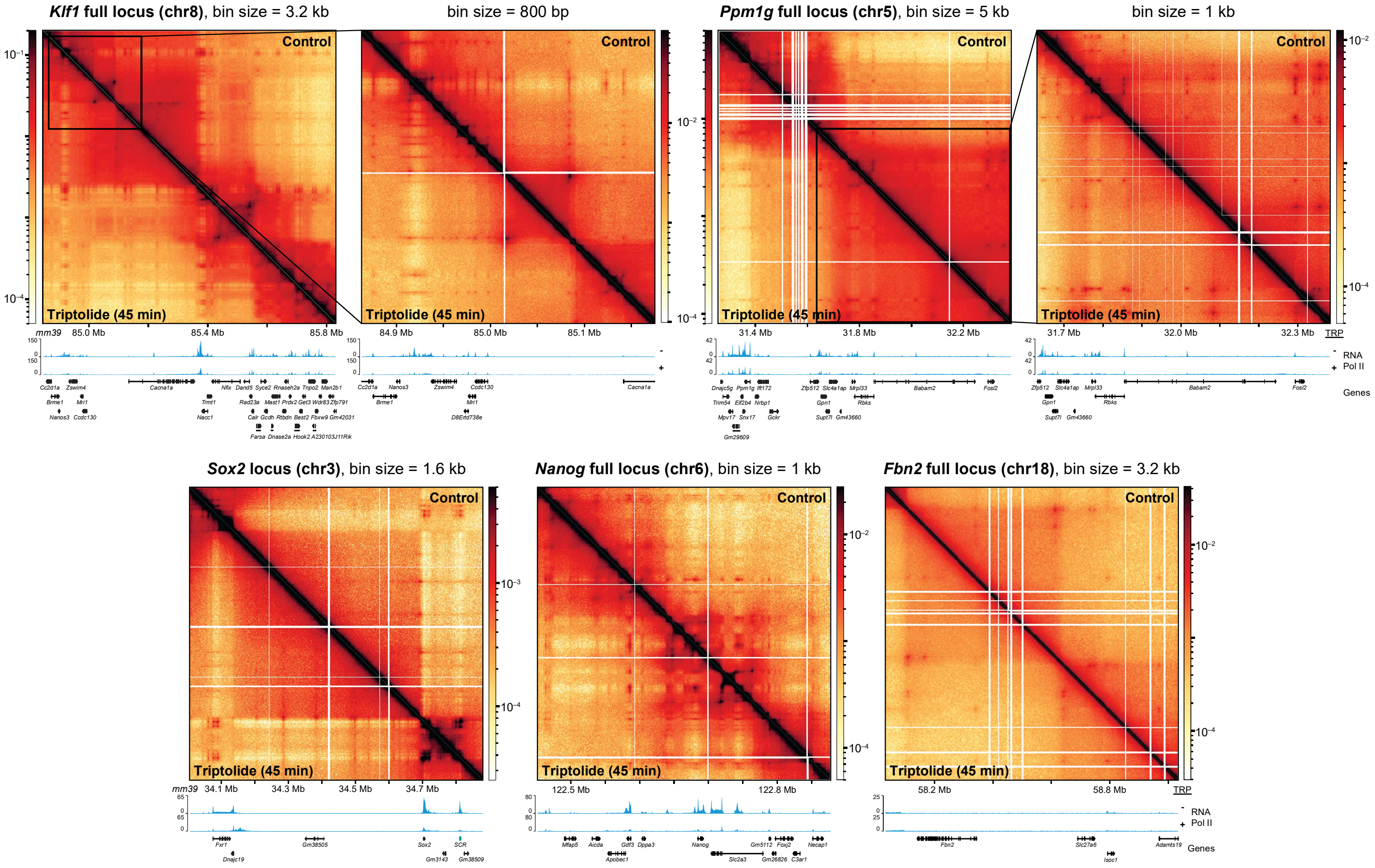


**Supplementary Figure 9. Inhibition of transcription does not significantly alter genome organization in Captured loci.** Contact maps comparing a WT control (above) and after 45 min triptolide treatment (1 µM, 45 minutes; below) are shown for the *Klf1*, *Ppm1g*, *Sox2*, *Nanog*, and *Fbn2* loci at resolutions spanning 800 bp – 5 kb in mESC WT cells. RNA Pol II ChIP data from Hsieh *et al.*[3] is shown below the maps before and after the triptolide treatment (1 µM, 45 minutes).

### Supplementary Table

| Sample | Description | GEO / ENCODE# | Figures | Reference |
|---|---|---|---|---|
| ChIP, CTCF | Architectural protein, loop extrusion factor | GSE90994 | 2, 3, S1, S3, S4, S5, S7 | Hansen AS, Pustova I, Cattoglio C, Tjian R et al. CTCF and cohesin regulate chromatin loop stability with distinct dynamics. Elife 2017 May 3;6. PMID: 28467304 |
| ChIP, SMC1A | Architectural protein, loop extrusion factor | GSE123636 | 2, 3, S1, S3, S4, S5, S7 | Hansen AS, Hsieh TS, Cattoglio C, Pustova I et al. Distinct Classes of Chromatin Loops Revealed by Deletion of an RNA-Binding Region in CTCF. Mol Cell 2019 Nov 7;76(3):395-411.e13. PMID: 31522987 |
| ChIP, H3K4me1 | Histone marker, enhancers | ENCFF282RLA | 2, 3, S3, S4, S5, S7 | ENCODE Project Consortium.. An integrated encyclopedia of DNA elements in the human genome. Nature 2012 Sep 6;489(7414):57-74. PMID: 22955616 |
| ChIP, H3K4me3 | Histone marker, active genes | ENCFF523UIR | 2, 3, S3, S4, S5, S7 | ENCODE Project Consortium.. An integrated encyclopedia of DNA elements in the human genome. Nature 2012 Sep 6;489(7414):57-74. PMID: 22955616 |
| ChIP, H3K36me3 | Histone marker, gene bodies | ENCFF848LEV | S7 | ENCODE Project Consortium.. An integrated encyclopedia of DNA elements in the human genome. Nature 2012 Sep 6;489(7414):57-74. PMID: 22955616 |
| ChIP, H3K27ac | Histone marker, enhancers | GSE90893 | 2, 3, S3, S4, S5, S7 | Chronis C, Fiziev P, Papp B, Butz S et al. Cooperative Binding of Transcription Factors Orchestrates Reprogramming. Cell 2017 Jan 26;168(3):442-459.e20. PMID: 28111071 |
| ChIP, P300 | Enhancer factor | GSE90893 | S7 | Chronis C, Fiziev P, Papp B, Butz S et al. Cooperative Binding of Transcription Factors Orchestrates Reprogramming. Cell 2017 Jan 26;168(3):442-459.e20. PMID: 28111071 |
| ChIP, MED1 | General transcription factor | GSE22562 | 2, 3, S3, S4, S5, S7 | Kagey MH, Newman JJ, Bilodeau S, Zhan Y et al. Mediator and cohesin connect gene expression and chromatin architecture. Nature 2010 Sep 23;467(7314):430-5. PMID: 20720539 |
| ChIP, NANOG | Pluripotency transcription factor | GSE71932 | S7 | Murakami K, Günesdogan U, Zylicz JJ, Tang WWC et al. NANOG alone induces germ cells in primed epiblast in vitro by activation of enhancers. Nature 2016 Jan 21;529(7586):403-407. PMID: 26751055 |
| ChIP, OCT4 | Pluripotency transcription factor | GSE90893 | S7 | Chronis C, Fiziev P, Papp B, Butz S et al. Cooperative Binding of Transcription Factors Orchestrates Reprogramming. Cell 2017 Jan 26;168(3):442-459.e20. PMID: 28111071 |
| ChIP, SOX2 | Pluripotency transcription factor | GSE90893 | S7 | Chronis C, Fiziev P, Papp B, Butz S et al. Cooperative Binding of Transcription Factors Orchestrates Reprogramming. Cell 2017 Jan 26;168(3):442-459.e20. PMID: 28111071 |
| ChIP, c-MYC | Pluripotency transcription factor | GSE90893 | S7 | Chronis C, Fiziev P, Papp B, Butz S et al. Cooperative Binding of Transcription Factors Orchestrates Reprogramming. Cell 2017 Jan 26;168(3):442-459.e20. PMID: 28111071 |
| ChIP, ESRRB | Enhancer factor | GSE90893 | S7 | Chronis C, Fiziev P, Papp B, Butz S et al. Cooperative Binding of Transcription Factors Orchestrates Reprogramming. Cell 2017 Jan 26;168(3):442-459.e20. PMID: 28111071 |
| ChIP, KLF4 | Pluripotency transcription factor | GSE90893 | S7 | Chronis C, Fiziev P, Papp B, Butz S et al. Cooperative Binding of Transcription Factors Orchestrates Reprogramming. Cell 2017 Jan 26;168(3):442-459.e20. PMID: 28111071 |
| ChIP, YY1 | Architectural protein, transcription factor | GSE99518 | S7 | Weintraub AS, Li CH, Zamudio AV, Sigova AA et al. YY1 Is a Structural Regulator of Enhancer-Promoter Loops. Cell 2017 Dec 14;171(7):1573-1588.e28. PMID: 29224777 |
| ChIP, H2AZ | Variant of H2A; found at active promoters | GSE51579 | S7 | Obri A, Ouararhni K, Papin C, Diebold ML et al. ANP32E is a histone chaperone that removes H2A.Z from chromatin. Nature 2014 Jan 30;505(7485):648-53. PMID: 24463511 |
| ChIP, RING1B | Repressive chromatin factor | GSE96107 | 2, 3, S3, S4, S5, S7 | Bonev B, Mendelson Cohen N, Szabo Q, Fritsch L et al. Multiscale 3D Genome Rewiring during Mouse Neural Development. Cell 2017 Oct 19;171(3):557-572.e24. PMID: 29053968 |
| ChIP, EZH2 | Repressive chromatin factor | GSE85717 | S7 | Juan AH, Wang S, Ko KD, Zare H et al. Roles of H3K27me2 and H3K27me3 Examined during Fate Specification of Embryonic Stem Cells. Cell Rep 2016 Oct 25;17(5):1369-1382. PMID: 27783950 |
| ChIP, H3K27me3 | Histone marker, repressive chromatin | GSE90893 | 2, 3, S3, S4, S5, S7 | Chronis C, Fiziev P, Papp B, Butz S et al. Cooperative Binding of Transcription Factors Orchestrates Reprogramming. Cell 2017 Jan 26;168(3):442-459.e20. PMID: 28111071 |
| ChIP, RNA Pol II | Transcription | GSE58019 | 2, 3, S3, S4, S5, S7 | Riising EM, Comet I, Leblanc B, Wu X et al. Gene silencing triggers polycomb repressive complex 2 recruitment to CpG islands genome wide. Mol Cell 2014 Aug 7;55(3):347-60. PMID: 24999238 |
| RNA-seq | Transcription | GSE123636 | 2, 3, S3, S4, S5, S7 | Hansen AS, Hsieh TS, Cattoglio C, Pustova I et al. Distinct Classes of Chromatin Loops Revealed by Deletion of an RNA-Binding Region in CTCF. Mol Cell 2019 Nov 7;76(3):395-411.e13. PMID: 31522987 |
| ATAC | Chromatin accessibility assay | GSE98390 | 2, 3, S3, S4, S5, S6, S7 | King HW, Fursova NA, Blackledge NP, Klose RJ. Polycomb repressive complex 1 shapes the nucleosome landscape but not accessibility at target genes. Genome Res 2018 Oct;28(10):1494-1507. PMID: 30154222 |
| ChIP, CTCF | Architectural protein, loop extrusion factor | GSE178982 | 4, S8 | Hsieh TS, Cattoglio C, Slobodyanyuk E, Hansen AS et al. Enhancer-Promoter Interactions and Transcription are Maintained Upon Acute Loss of CTCF, Cohesin, WAPL, and YY1. bioRxiv 2021 Jul 14:452-365. doi: https://doi.org/10.1101/2021.07.14.452365 |
| ChIP, RAD21 | Architectural protein, loop extrusion factor | GSE178982 | 4, S8 | Hsieh TS, Cattoglio C, Slobodyanyuk E, Hansen AS et al. Enhancer-Promoter Interactions and Transcription are Maintained Upon Acute Loss of CTCF, Cohesin, WAPL, and YY1. bioRxiv 2021 Jul 14:452-365. doi: https://doi.org/10.1101/2021.07.14.452365 |
| ChIP, YY1 | Architectural protein, transcription factor | GSE178982 | 4, S8 | Hsieh TS, Cattoglio C, Slobodyanyuk E, Hansen AS et al. Enhancer-Promoter Interactions and Transcription are Maintained Upon Acute Loss of CTCF, Cohesin, WAPL, and YY1. bioRxiv 2021 Jul 14:452-365. doi: https://doi.org/10.1101/2021.07.14.452365 |
| RNA-seq | Transcription | GSE178982 | 4, S8 | Hsieh TS, Cattoglio C, Slobodyanyuk E, Hansen AS et al. Enhancer-Promoter Interactions and Transcription are Maintained Upon Acute Loss of CTCF, Cohesin, WAPL, and YY1. bioRxiv 2021 Jul 14:452-365. doi: https://doi.org/10.1101/2021.07.14.452365 |
| ChIP, RNA Poll II | Transcription | GSE130275 | 4, S9 | Hsieh TS, Cattoglio C, Slobodyanyuk E, Hansen AS et al. Resolving the 3D Landscape of Transcription-Linked Mammalian Chromatin Folding. Mol Cell 2020 May 7;78(3):539-553.e8. PMID: 32213323 |

#### Materials and Methods
##### *Experimental Procedure*

**Overview of the Region Capture Micro-C experiment**

Region Capture Micro-C (RCMC) was developed by merging Micro-C[3] with tiled Capture of a locus[11,12]. An overview of the RCMC protocol is provided below, and a detailed protocol is attached as a supplementary protocol. The data generated in this manuscript comes from the merger of three RCMC replicates, each of which were generated by pooling five 5M cell pellets for each of the four tested conditions (wild-type, transcriptional inhibition, cohesin depletion, and a DMSO-treated control); reaction volumes for both 1M and 5M cell samples are provided in the attached protocol. RCMC replicates are technical replicates; 125-200M cells were harvested (cultured, crosslinked, aliquoted, and snap-frozen) for each tested condition, after which downstream RCMC steps (Micro-C and tiled Capture) were replicated several times for each snap-frozen aliquot. However, RCMC data for untreated wild-type mESC and DMSO-vehicle treated F1M RAD21-mAID-BFP-V5 JM8.N4 mESCs represent biological replicates in the sense of reporting general mESC 3D genome structure for two independent clones without perturbation.

**Cell culture**

Mouse embryonic stem cells (JM8.N4 mESCs[13]) were cultured on plates coated with 0.1% gelatin solution (Sigma-Aldrich #G1890) under feeder-free conditions in medium consisting of KnockOut DMEM (ThermoFisher #10829-018) with 15% FBS (HyClone, SH30396.03, Lot. No. AE28209315) and 1000U/mL LIF (home-made[14]), 1 mM MEM Non-Essential Amino Acid Solution (ThermoFisher #11140-050), 2 mM GlutaMAX (ThermoFisher #35050061), 100 µg/ml Penicillin-Streptomycin (ThermoFisher #15140-122), and 0.1 mM 2-mercapoethanol (ThermoFisher #31350010) supplemented with 2i, 10 µM MEK inhibitor (Tocris #PD0325901) and 3 µM GSK inhibitor (Sigma-Aldrich #SML1046). mESCs were fed daily by replacing half of the medium and passaged every two days with TrypLE Express Enzyme (ThermoFisher #12605036). One day prior to treatment and harvesting, cells were swapped to medium as described above without 2i.

**Depletion of cohesin**

Depletion of cohesin was achieved using indole-3-acetic acid (IAA) treatment as previously described[9,10]. A 250 mM IAA (BioAcademia #30-003-10) stock was prepared by dissolving the drug in DMSO. F1M RAD21-mAID-BFP-V5 JM8.N4 mESCs[9] were grown to ~80% confluency in medium as described above, with a swap to 2i-free medium 24 hours prior to treatment. Cells were washed once with PBS and fed fresh 2i-free medium containing either only DMSO (untreated control) or 500 µM IAA (cohesin depleted), incubated for 3 hours and then harvested.

**Inhibition of transcription**

Inhibition of RNA Pol II activity was achieved using triptolide treatment as previously described[3]. A 1 mM triptolide (Sigma-Aldrich #T3652) stock was prepared by dissolving the drug in DMSO. Wild-type JM8.N4 mESCs were grown to ~80% confluency in medium as described above, with a swap to 2i-free medium 24 hours prior to treatment. Cells were washed once with PBS and fed fresh 2i-free medium containing 1 µM triptolide, incubated for 45 min, and then harvested.

**Crosslinking**

Cells were doubly crosslinked to fix protein-protein and protein-DNA interactions using DSG (disuccinimidyl glutarate, 7.7Å) (ThermoFisher #20593) and formaldehyde (ThermoFisher #28906), respectively. Crosslinking medium was prepared by diluting freshly made DSG stock solution (300 mM DSG in DMSO) to 3 mM in 1X PBS (ThermoFisher #10010031). Trypsinized cells were resuspended to single cells, counted, washed in PBS, and then resuspended in crosslinking medium at a concentration of 1M cells per mL. The crosslinking reaction was gently mixed at room temperature for 35 minutes, after which formaldehyde was added to a final concentration of 1%. The double crosslinking reaction was mixed at room temperature for an additional 10 minutes before quenching with Tris buffer pH = 7.5 (K-D Medical #RGE-3370) at a final concentration of 0.375 M. Crosslinked cells were washed twice with 1X PBS, re-counted to quantify any sample loss during fixation, and then partitioned into 5M cell aliquots that were pelleted and snap-frozen in liquid nitrogen for storage at –80°C.

**Micrococcal nuclease (MNase) titration**

Digesting the crosslinked genome to the nucleosome-sized fragments (150-200 bp) necessary to capture nucleosome-resolution DNA contacts requires a titration to identify the ideal MNase digestion concentration and reaction conditions. Accordingly, MNase titrations were performed for each batch of crosslinked cells before performing the RCMC protocol. The titration involved MNase digestion on 1M or 5M cell samples varying MNase concentrations, reversal of crosslinks, DNA purification, and gel-based separation to visualize the distribution of fragment sizes (see corresponding sections below). Ideal digestion concentrations were identified by samples digested to primarily (~80%) mononucleosomal fragments (150-200 bp), few (~15-20%) dinucleosomal fragments (250-350 bp), and a faint but visible band (<5%) of trinucleosomal fragments (400-500 bp) (**Fig. S1a**).

**Micrococcal nuclease digestion**

Cell membranes were solubilized to extract intact nuclei by resuspending crosslinked 5M cell pellets in Micro-C Buffer #1 (MB#1; 50 mM NaCl, 10 mM Tris-HCl pH = 7.5, 5 mM MgCl2, 1M CaCl2, 0.2% NP-40 Alternative (Millipore Sigma #492018), 1x Protease Inhibitor Cocktail (Sigma-Aldrich #5056489001)) at 1M cells per 100 µL for 20 min on ice. Following an MB#1 wash, samples were resuspended in 100 µL MB#1 and the amount of 20 U/µL MNase (Worthington Biochem #LS004798) determined by MNase titration was added. This digestion reaction was mixed at 37°C for 20 min on a thermomixer before being quenched with 4 mM EGTA (bioWORLD #40520008) and heat inactivated at 65°C for 10 min. Digested nuclei were washed twice with ice-cold Micro-C Buffer #2 (50 mM NaCl, 10 mM Tris-HCl pH = 7.5, 10 mM MgCl2, 100 µg/mL BSA (Sigma-Aldrich #B8667)).

**End repair and labeling**

To generate blunt ends on digested DNA fragments prior to proximity ligation and add biotinylated nucleotides, a series of enzymatic processing steps were performed. First, to catalyze the addition of 5'-phosphate groups and the removal of 3'-phosphate groups, digested samples generated from 5M cell inputs were incubated in end-repair reactions (50 U T4 Polynucleotide Kinase (New England BioLabs #M0201), 50 mM NaCl, 10 mM Tris-HCl pH = 7.5, 10 mM MgCl2, 100 ug/mL BSA, 2 mM ATP (ThermoFisher #R1441), 5 mM DTT (Sigma-Aldrich #10197777001), in water) at 37°C for 15 min while mixing. To create 5' fragment overhangs for end-blunting and labelling, 50 U of DNA Polymerase I Klenow Fragment (New England BioLabs #M0210) was added to the reaction and incubated at 37°C for 15 min while mixing. Next, a mixture of dNTPs in end-labelling buffer (66 µM each of dTTP (Jena Bioscience #NU-1004), dGTP (Jena Bioscience #NU-1003), biotin-dATP (Jena Bioscience #NU-835-BIO14), and biotin-CTP (Jena Bioscience #NU-809-BIOX), 1X T4 DNA Ligase Buffer, 100 µg/mL BSA, in water) was added to the reaction. This reaction was incubated at room temperature for 45 min with interval mixing before being quenched by 30 mM EDTA (Invitrogen #15575020) and heat inactivated at 65°C for 20 min. Finally, end-blunted and biotin-labeled nuclei were washed once with Micro-C Buffer #3 (50 mM Tris-HCl pH = 7.5, 10 mM MgCl2, 100 µg/mL BSA).

**Proximity ligation and removal of unligated biotin**

Proximity ligation was performed by incubating labeled chromatin in a ligation reaction (10,000 U T4 DNA Ligase (New England BioLabs #M0202), 1X T4 DNA Ligase Buffer, 100 µg/mL BSA, in

500 µL water) at room temperature for at least 2.5 hours with gentle mixing. To remove biotinylated dNTPs from all unligated fragment ends, samples were digested by 1,000 U of Exonuclease III (New England BioLabs #M0206) in reaction buffer (1X NEBuffer #1 in water) at 37°C for 15 min with interval mixing.

**DNA purification and size-selection**

In order to prepare ligated DNA for library generation, DNA was reverse crosslinked and proteins and RNA were digested by adding 1% SDS (Sigma-Aldrich #L3771), 2 mg/mL Proteinase K (Viagen Biotech #501-PK), 250 mM NaCl, and 100 µg/mL RNaseA (ThermoFisher #EN0531) to the samples and incubating at 65°C overnight. DNA was extracted using phenol:chloroform:isoamylic alcohol (PCI) (Sigma-Aldrich #P2069) in a 1:1 volumetric ratio using 5PRIME Phase Lock Gel Light tubes (Quantabio #2302820). The aqueous phase was further purified using the Zymo DNA Clean & Concentrator kit (Zymo Research #D4034) according to the kit manual.

Dinucleosome-sized DNA fragments (250-350 bp) were isolated by extraction from a 1% agarose gel (VWR #97062) (**Fig. S1b**). Gel extracts were purified using the Zymo Gel Purification kit (Zymo Research #D4008), and samples were quantified by Qubit 1X dsDNA High Sensitivity Assay (Invitrogen #Q33231). Sample ends were polished and blunted again using the End-It enzyme reaction (Lucigen #ER81050) at 25°C for 45 minutes, followed by reaction inactivation at 65°C for 10 min.

Ligated DNA contact fragments were isolated by pulling down biotin-bound fragments using Dynabeads MyOne Streptavidin T1 (Invitrogen #65601). DNA samples were bound to beads in a Binding and Wash Buffer (1 M NaCl, 5 mM Tris-HCl pH = 7.5, 500 µM EDTA, 0.1% Tween-20 (Sigma-Aldrich #P8074)) at room temperature for at least 30 minutes with mixing. After two washes with the Binding and Wash Buffer, the bead-bound samples were washed once with 10 mM Tris-HCl pH = 7.5 prior to library prep.

**Library preparation**

Illumina library preparation was performed using the NEBNext Ultra II kit (New England BioLabs #E7645) to end-repair, A-tail, and adaptor ligate the bead-bound samples. All steps were performed as directed by the manual, except that incubations included interval shaking (1 minute on, 3 minutes off) at 1000 rpm. Sample washes were performed using Binding and Wash Buffer and 10 mM Tris-HCl pH = 7.5 washes. To determine the minimum number of PCR cycles to meet input material guidelines for capture or sequencing, a test library amplification was performed with 5% or less of the prepped library to quantify the yield. The test PCR reaction mixture was run on an agarose gel and yield quantified using image quantification software Image Studio Lite (LI-COR Biosciences). 10 or fewer PCR cycles to meet Capture input requirements is optimal to reduce PCR duplicates, and the RCMC replicates in this manuscript used 7-8 PCR cycles for final library amplification. All library amplifications were done using sequencing indices from the NEB Multiplex Oligos for Illumina Primer Set 1 (New England BioLabs #E7335) and the KAPA HiFi HotStart ReadyMix enzyme (Roche #07958927001). Following library amplification, the T1 Dynabeads containing the original bead-bound samples were removed and the amplified libraries were purified to remove adaptor dimers, primers, and contaminants using AmPure XP beads (Beckman Coulter #A63880). Purified libraries were quantified via Fragment Analyzer and qPCR at the MIT BioMicro Center to determine library concentrations for pooling prior to Capture.

**Capture probe design**

Target loci of interest were identified based on genomic features or enhancer-promoter relationships of interest. *Klf1* and *Ppm1g* were selected as gene-rich loci, *Fbn2* was selected as a gene-poor control with a well-established CTCF- and cohesin-mediated loop[9], and *Sox2* and *Nanog* were later selected as loci for comparing RCMC against TMCC (**Fig. S3**). Using the UCSC Genome Browser and HiGlass visualization of existing mESC 3C datasets, locus bounds were selected to include visible local structures and genomic features in roughly 1 Mb-sized regions. Once loci had been selected, 80-mer probes were designed to tile end-to-end without overlap across the Capture loci through Twist Bioscience (**Fig. S1c**). Probes with high predicted likelihoods of off-target pulldown (e.g. such as those in high-repeat regions) were masked and removed from the probe tiling, and probe coverage was double-checked to ensure the inclusion of key genomic features (e.g. all promoters and CTCF sites in the locus) before finalization. Probe panels were synthesized and purchased as Custom Target Enrichment Panels from Twist Bioscience.

**Capture of target loci**

Capture was performed in accordance with Twist Bioscience's Standard Hybridization Target Enrichment Protocol. Briefly, pooled sample libraries were dried and mixed with Hybridization Mix (Twist Bioscience #104178), Custom Panels (Twist Bioscience #101001), and Universal Blockers (Twist Bioscience #100578), as well as Mouse Cot-1 DNA (Invitrogen #18440016). The library pool was hybridized to the biotinylated probe panel overnight, after which streptavidin beads (Twist Bioscience #100983) were used to pull down probes with hybridized ligated fragments and then washed (Twist Bioscience #104178) to remove unbound fragments. Another round of PCR amplified the target-enriched library using the Equinox Library Amplification Mix (Twist Bioscience #104178), with including a test PCR (as described above) to identify the number of amplification cycles necessary to meet sequencing requirements. With 2-4 µg of input library for Capture, the RCMC samples generated in this manuscript needed 6 cycles of post-Capture PCR amplification. Following PCR amplification, the Captured library was purified (Twist Bioscience #100983) and then quantified via both Fragment Analyzer and qPCR at the MIT BioMicro Center in preparation for sequencing submission.

Three technical replicates of the pre-Capture Micro-C library were generated, after which each replicate was simultaneously Captured for the *Klf1*, *Ppm1g*, and *Fbn2* loci. After the publication of TMCC[5], additional probes for the *Sox2* and *Nanog* loci were designed and a single additional Capture experiment was conducted pooling all three pre-Capture Micro-C libraries for simultaneous *Sox2* and *Nanog* Capture.

**Sequencing**

Following qPCR quantification, post-Capture libraries across the four samples (wild-type, transcriptionally inhibited, cohesin depleted, and DMSO-treated control) were pooled in a 1:1 molar ratio. Pooled libraries were sequenced by paired-end 2x50 cycle sequencing kits with Illumina NovaSeq SP or S1 flow cells on a NovaSeq 6000 system by the Broad Institute of MIT and Harvard's Walk-Up Sequencing services. Basecalls for NovaSeq output were performed using bcl2fastq v2.20.0.422.

***Data Analysis***

**Mapping and normalizing RCMC**

RCMC paired-end reads generated by the Illumina NovaSeq sequencers were downloaded as .fastq files for each sample, pair mate, and flow cell lane. Read quality was verified using FastQC (v0.11.9). Paired end reads were aligned to the UCSC mm39 genome using bowtie2 (v2.3.5.1) with --local --reorder --very-sensitive-local. Aligned paired end reads were then parsed with pairtools (v0.3.0) parse with --add-columns mapq --walks-policy mask --min-mapq 2. Parsed reads were filtered for PCR duplicates and unmapped/mutiple mapping reads with pairtools dedup with --max-mismatch 1. Remaining reads were indexed (pairix v0.3.7) and filtered (pairtools select) to retain only those reads where both read mates lie in a locus of interest. These filtered reads were subsequently converted to .cool format using cooler (v0.8.11) cload pairs, creating binned read counts across the genome for 50 bp bins. Finally, .cool files were converted to the .mcool format with cooler zoomify including the --balance option, compiling read counts for bins from 50 bp up to 10 Mb in size.

Contact matrices were balanced using iterative correction and eigendecomposition (ICE)[1], which normalizes all rows and columns of a contact matrix sum to the same value. Applying ICE balancing to all mapped reads generated subpar normalization and generated an artifact where "stripes" containing no Capture probe coverage appeared to have greater contact densities than adjacent probe-covered regions (**Fig. S1e**). ICE balancing to .mcool files containing data only within Captured regions of interest (ROIs) did not result in these artifacts, and was therefore used in for all RCMC data in this study. The success of ICE balancing applied to these ROI-only .mcools was evaluated against published whole-genome Hi-C[2] and Micro-C[3] datasets in mESCs (**Fig. S6a**). The sum of each row of each of the RCMC, Micro-C, and Hi-C balanced contact matrices at 250 bp resolution within Capture ROIs was calculated, plotted as a histogram distribution of row sums, and verified to match the distribution of column sums. The subset of RCMC rows containing microcompartment anchors was also plotted to confirm that they match the distribution of row sums across the whole locus, ruling out that microcompartments are an artifact of incomplete ICE-normalization[1].

**Visualizing RCMC**

RCMC contact maps were visualized alongside genomic annotations, published ChIP-seq, RNA-seq, and ATAC-seq datasets using the HiGlass[15] browser (http://higlass.io/) and software (v0.8.0). Contact maps shown in figures were generated using cooltools (v0.5.0) (https://cooltools.readthedocs.io/). Genomic tracks (i.e. ChIP-seq, RNA-seq, and ATAC-seq) and gene annotations for manuscript figures were generated using CoolBox[16] (v0.3.3). In generating our genomic tracks, we analyzed 27 public datasets (Supplementary Table 1) using processed bigWig files which were CrossMapped[17] (v0.6.1) (http://crossmap.sourceforge.net/) to the mm39 reference genome. Tracks were visualized using the Integrative Genomics Viewer (IGV)[18] (v2.10.3) to scale tracks by identifying local maxima and minimizing noise.

**Comparing data across methods**

Mapped sequencing reads were filtered using pairtools select to quantify read counts according to chosen evaluation criteria (**Fig. 1b, Fig. S1d, 2c**). Filtering was performed identically across the RCMC, TMCC, Micro-C, and Hi-C datasets on .pairs files containing mm39-mapped reads. RCMC .pairs files were generated as described above, while mm10-aligned .pairs files containing all unique reads were downloaded for Hi-C (GSE96107) and Micro-C (GSE130275) and CrossMapped to the mm39 genome. TMCC .pairs files were generated through two methods. For TMCC contact map generation and all analyses using unique contacts (**Fig. 1b, 2c, Fig. S2a-c, 4b**), curated lists of unique contacts (in mm10) were downloaded for wild-type TMCC data (GSE181694), converted to a .pairs data format, and CrossMapped to the mm39 genome. For quantifications of total mapped reads (**Fig. S2c, downsampling in Fig. 2c and Fig. S4b**), raw sequencing data files were downloaded for TMCC and aligned to the mm39 genome in a similar manner to RCMC.

Quantifications of read coverage across bins (**Fig. 1b, Fig. S2b**) were calculated in Python using cooler to load unbalanced 100 bp resolution .cool files into memory as matrices. These matrices were then iterated through to determine the fractions of bins containing at least one read at different contact distances.

Genome-wide equivalents for RCMC data were calculated by extrapolating the number of unique contacts mapped to a Capture locus to a region the size of the entire mouse genome. This approach assumes homogeneous read coverage throughout the genome; in reality, however, read coverage is unevenly distributed between regions depending on the specific region and which 3C method is used. Specifically, both genome-wide Hi-C[2] (3.3B total unique) and Micro-C[3] (2.64B total unique) also had higher coverage at the *Klf1* region than the genome-wide average. As such, compared to Hi-C[2] at *Klf1*, RCMC captured ~38-fold more unique contacts. Similarly, compared to genome-wide Micro-C[3] at *Klf1*, RCMC captured ~12-fold more unique contacts.

**Contact decaying curve analysis**

Contact decay curves were generated by plotting contact probability against genomic separation using cooltools (**Fig. S2a, 8b**). Balanced and smoothed curves were generated using contact matrices across each chromosome binned to 50 bp resolution. RCMC and TMCC curves were truncated at 1 Mb genomic separation due to noise at larger genomic separations (the largest Captured locus for each method is between 1-2 Mb in size).

**Replicate reproducibility analysis**

The reproducibility of RCMC technical replicates (**Fig. S1g, 8c**) was evaluated using HiCRep[4] (v1.12.2) for contact maps at 5 kb resolution, with parameters lbr = 0 and ubr = 5000000. Reproducibility scores were calculated for each region independently for all replicates using the optimal h-value determined from a single replicate.

**Downsampling**

Tiled-Micro-Capture-C (TMCC) was downsampled using pairtools sample to randomly select a subset of the pairs in the locus of interest. To ensure parity in comparison, .pairs files containing all mapped reads (all unique and duplicate reads with both mates in the viewpoint) were first generated for viewpoints of interest in the *Sox2* (**Fig. S4b**) and *Nanog* (**Fig. 2c**) loci. These files yielded the total number of mapped reads necessary to generate the density of information present in the viewpoint, and a downsampling ratio was calculated to match the total number of mapped TMCC reads in the viewpoint with RCMC. After a randomly downsampled TMCC .pairs file was generated, the unique reads were extracted and used to generate a .mcool for visualization purposes.

**Chromatin contact analysis**

Chromatin loops were called on WT Capture ROI-only RCMC data using Mustache[6] (v1.2.4) (https://github.com/ay-lab/mustache) at 0.25, 0.5, 1, 2, 5, and 10 kb data resolutions with sparsity thresholds of 0.7 and q-value thresholds of 0.1 (**Fig. S5c**). Loop-calling was also tested using Chromosight[19] (v1.6.1) (https://github.com/koszullab/chromosight) and SIP[20] (v1.6.1) (https://github.com/PouletAxel/SIP), with Mustache producing similar or greater numbers of loop calls. Finer resolutions of loop calling called more microcompartmental contacts, but still missed many contacts while also increasingly misidentifying stripes as loops, overlapping loops, and clustering calls at short contact distances off of the diagonal.

Manual contact-calling  was performed in an attempt to minimize these artifacts in microcompartment analysis. We defined contacts as punctate foci of interaction (i.e. “dots”), visibly discernible as being enriched relative to their local background. We did not identify diffuse and overly faint interactions, homogeneously enriched stripes, and short-range contacts just off of the diagonal (i.e. under ~5 kb contact distance) as contacts. Contacts were called on ICE-balanced WT Capture ROI-only RCMC data at 250 bp resolution across the full *Klf1* and *Ppm1g* loci using the HiGlass browser interface. Scale bar limits were dynamically modulated to minimize background and clearly distinguish focal enrichment. A total of 1093 focal contacts (loops) spanning 132 contact anchors were manually annotated across the two loci (**Fig. S5d**). Called contacts ranged in length from 4.6 to 1,018 kb between anchors, with a strong majority of contacts laying in the *Klf1* locus.

**Compartmentalization analysis**

Compartments were called by applying eigendecomposition to the WT RCMC contact matrix using cooltools (**Fig. S5c**). Capture ROI-only RCMC data was first normalized to remove the distance-dependent effect of contact frequency. Eigendecomposition was then performed, with GC content serving as a correlate for orienting eigenvectors to indicate A- (gene-rich or active chromatin) or B- (gene-poor or inactive chromatin) compartments. Finally, eigenvectors were binarized and visualized as BED tracks using CoolBox. Compartment calling was performed at 0.5, 1, 5, and 50 kb data resolutions, with all producing similar output (calls at 1 kb resolution are shown for clarity's sake) or outright failing to call compartments.

**Microcompartment anchor classification**

To classify microcompartments anchors as promoter, enhancer, or CTCF and cohesin-bound (**Fig. 3e**), microcompartment anchor locations were compared with the corresponding chromatin features as follows. Promoter regions were defined using all TSS locations in the mm39 UCSC RefGene annotation[21] ±2 kb. Enhancers or CTCF and cohesin-bound sites were defined based on overlap of H3K4me1 (ENCFF282RLA) and H3K27ac (GSE90893), or CTCF (GSE90994) and SMC1A (GSE123636), respectively. For all datasets, bigwig files were converted to bedgraph files using UCSC bigWigToBedGraph (v377)[22], followed by peak calling using MACS2 bdgpeakcall[23] (v2.2.7.1). For CTCF, called peaks were then overlapped with CTCF sites identified using FIMO (v5.4.1): first, fasta-get-markov was used to generate a background model using the mm39 genome assembly, then motifs were identified using –max-stored-scores 50000000 –thresh 1e-3. Finally, locations of motifs were overlapped with peaks identified in ChIP-seq data, and only the motif with the highest score for each peak was maintained. The peaks (H3K4me1, H3K27ac, SMC1A) or identified sites (CTCF) were then overlapped using bedtools intersect (v2.30.0) to give enhancers or CTCF and cohesin-bound regions. Anchors of microcompartments ±1 kb were then overlapped with each of the three features to classify them as promoter, enhancer, or CTCF and cohesin-bound. Regions which were classified as both promoter and enhancer regions were treated as promoter-only due to the inability to distinguish H3K4me1/H3K27ac overlap at enhancers and promoters. Anchors overlapping none of these three features were classified as "Other". Anchor classifications were then considered combinatorially to classify microcompartment interactions between regions (**Fig. 3g**).

Anchor overlap with ATAC-seq (GSE98390) peaks and vice versa was determined in the same way as for ChIP-seq data, after limiting ATAC-seq peaks to only those falling within the captured *Ppm1g* and *Klf1* regions, to identify sets of regions which did or did not overlap (**Fig. S6d**).

The number of interactions formed by each anchor and the lengths of the interactions they form (**Fig. 3c, d, f**) was determined and visualized in R[24] (v4.1.2) using the GenomicRanges[25] (v1.46.1) and ggplot2[26] (v3.3.6) packages.

**Heatmap and metaplot generation**

Heatmaps and metaplots were generated for microcompartments and ATAC-seq peaks using deeptools[27] (v3.5.1) computeMatrix followed by plotHeatmap in a region ±1 kb around the center of each site for genomics data listed in Supplementary Table 1 (**Fig. S6b, 7**). Regions were sorted in all cases according to decreasing ATAC-seq signal.

**Pile-up analysis**

Aggregate peak analysis of microcompartmental contacts (**Fig. 4**) was performed using cooltools[7,8]. Plots for all contacts of a given classification (e.g. all wild-type Enhancer-Promoter microcompartments) were generated and averaged for a 20 kb window centered on the contact at 250 bp resolution. Background-normalized dot enrichment values were calculated using 1250x1250 bp (5x5 pixel) boxes; the dot's average intensity values were determined by a box in the center of the viewing window whereas the background's average intensity values were determined by boxes in the top left and bottom right corners of the viewing window (chosen for their equivalent contact distances as the dot).

**Data and code availability**

The raw and processed RCMC data reported in this manuscript are available at GEO: GSE207225.

### Supplementary References


1. Imakaev, M. *et al.* Iterative correction of Hi-C data reveals hallmarks of chromosome organization. *Nat. Methods* **9**, 999–1003 (2012).
2. Bonev, B. *et al.* Multiscale 3D Genome Rewiring during Mouse Neural Development. *Cell* **171**, 557-572.e24 (2017).
3. Hsieh, T.-H. S. *et al.* Resolving the 3D Landscape of Transcription-Linked Mammalian Chromatin Folding. *Mol. Cell* (2020). doi:https://doi.org/10.1016/j.molcel.2020.03.002
4. Yang, T. *et al.* HiCRep: assessing the reproducibility of Hi-C data using a stratum-adjusted correlation coefficient. *Genome Res.* **27**, 1939–1949 (2017).
5. Aljahani, A. *et al.* Analysis of sub-kilobase chromatin topology reveals nano-scale regulatory interactions with variable dependence on cohesin and CTCF. *Nat. Commun.* **13**, 2139 (2022).
6. Roayaei Ardakany, A. *et al.* Mustache: multi-scale detection of chromatin loops from Hi-C and Micro-C maps using scale-space representation. *Genome Biol.* **21**, 256 (2020).
7. Sergey Venev, Nezar Abdennur, Anton Goloborodko, Ilya Flyamer, Geoffrey Fudenberg, Johannes Nuebler, Aleksandra Galitsyna, Betul Akgol, Sameer Abraham, Peter Kerpedjiev, & M. I. open2c/cooltools: v0.4.1 (v0.4.1). *Zenodo* https://doi.org/10.5281/zenodo.5214125 (2021). doi:https://doi.org/10.5281/zenodo.5214125
8. Abdennur, N. *et al.* Cooler: scalable storage for Hi-C data and other genomically labeled arrays. *Bioinformatics* **36**, 311–316 (2020).
9. Gabriele, M. *et al.* Dynamics of CTCF- and cohesin-mediated chromatin looping revealed by live-cell imaging. *Science (80-. ).* **376**, 496–501 (2022).
10. Hsieh, T.-H. S. *et al.* Enhancer-promoter interactions and transcription are maintained upon acute loss of CTCF, cohesin, WAPL, and YY1. *bioRxiv* 2021.07.14.452365 (2021). doi:10.1101/2021.07.14.452365
11. Jäger, R. *et al.* Capture Hi-C identifies the chromatin interactome of colorectal cancer risk loci. *Nat. Commun.* **6**, 6178 (2015).
12. Oudelaar, A. M. *et al.* Dynamics of the 4D genome during in vivo lineage specification and differentiation. *Nat. Commun.* **11**, 2722 (2020).
13. Pettitt, S. J. *et al.* Agouti C57BL/6N embryonic stem cells for mouse genetic resources. *Nat. Methods* **6**, 493–495 (2009).
14. Hansen, A. S. *et al.* CTCF and cohesin regulate chromatin loop stability with distinct dynamics. *Elife* **6**, e25776 (2017).
15. Kerpedjiev, P. *et al.* HiGlass: web-based visual exploration and analysis of genome interaction maps. *Genome Biol.* **19**, 125 (2018).
16. Xu, W. *et al.* CoolBox: a flexible toolkit for visual analysis of genomics data. *BMC Bioinformatics* **22**, 489 (2021).
17. Zhao, H. *et al.* CrossMap: a versatile tool for coordinate conversion between genome assemblies. *Bioinformatics* **30**, 1006–1007 (2014).
18. Robinson, J. T. *et al.* Integrative genomics viewer. *Nat. Biotechnol.* **29**, 24–26 (2011).
19. Matthey-Doret, C. *et al.* Computer vision for pattern detection in chromosome contact maps. *Nat. Commun.* **11**, 5795 (2020).
20. Rowley, M. J. *et al.* Analysis of Hi-C data using SIP effectively identifies loops in organisms from C. elegans to mammals. *Genome Res.* (2020). doi:10.1101/gr.257832.119
21. Lee, B. T. *et al.* The UCSC Genome Browser database: 2022 update. *Nucleic Acids Res.* **50**, D1115–D1122 (2022).
22. Kent, W. J. *et al.* BigWig and BigBed: enabling browsing of large distributed datasets. *Bioinformatics* **26**, 2204–2207 (2010).
23. Zhang, Y. *et al.* Model-based Analysis of ChIP-Seq (MACS). *Genome Biol.* **9**, R137 (2008).
24. (2021), R. C. T. R: A language and environment for statistical computing. R Foundation for Statistical Computing. *https://www.R-project.org/* (2021).
25. Lawrence, M. *et al.* Software for Computing and Annotating Genomic Ranges. *PLOS Comput. Biol.* **9**, e1003118 (2013).
26. Wickham, H. Data analysis. ggplot2. in *Springer* 189–201 (2016).
27. Ramírez, F. *et al.* deepTools2: a next generation web server for deep-sequencing data analysis. *Nucleic Acids Res.* **44**, W160–W165 (2016).

### **Region Capture Micro-C (RCMC)** – Protocol Overview

RCMC is a chromosome conformation capture assay that merges Micro-C with tiled Capture of regions of interest, allowing deep mapping of 3D genome structure with relatively shallow sequencing. A diagram (below) details the RCMC workflow.

The RCMC protocol contains **3 sub-protocols**:


1. **RCMC Protocol, Pre-Capture** ........................................................................................ **pp. 2-14**


   - This is the bulk of the RCMC workflow and covers all experimental steps from cell culture through Micro-C library prep. Plan for this to take 3-5 days.


2. **MNase Titration Protocol** ........................................................................................ **pp. 15-19**


   - The RCMC workflow relies on digesting chromatin to primarily mononucleosomally-sized (e.g. 150-200 bp) fragments. Overdigestion of chromatin leads to short fragments and poor proximity ligation, whereas underdigestion risks mapping fewer contacts than possible.
   - Digestion time and conditions may vary from cell-type to cell-type and can vary by fixation; accordingly, it is advisable to do an MNase titration to determine the ideal MNase digestion amounts and conditions before proceeding with the rest of the protocol.
   - Conduct an MNase titration for each new batch of crosslinked cells. Plan for this to take 2 days.

3. **Capture Protocol** (*Twist Bioscience*) ............................... available on Twist's website, linked here
   - Capture is performed following the Standard Hybridization Target Enrichment Protocol provided by Twist Bioscience. The RCMC workflow follows it nearly verbatim, with deviations noted in the RCMC Protocol, Pre-Capture document. Plan for this to take 2 days.
   - Designing and ordering Capture probes can have a lag time of a several weeks, so do this proactively.

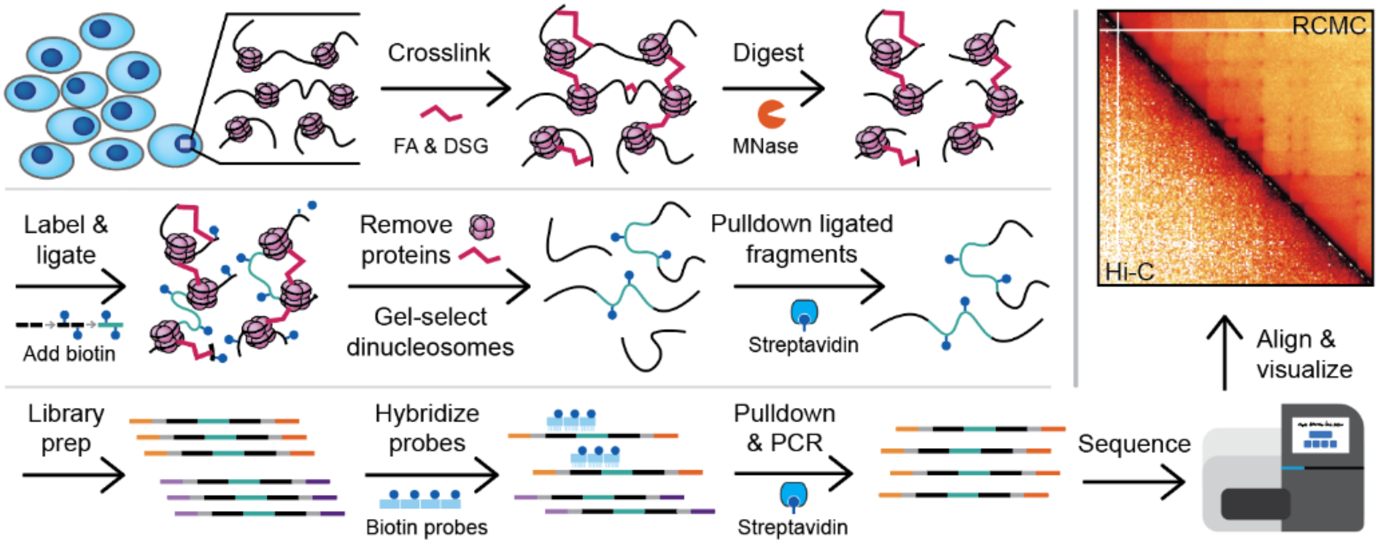


**Overview of the Region Capture Micro-C (RCMC) protocol.** Cells are chemically fixed, nuclei are digested with micrococcal nuclease (MNase), and fragments are biotinylated, proximity ligated, dinucleosomes gel extracted and purified, library prepped, PCR amplified, and region-enriched to create a sequencing library. After sequencing, mapping, and normalization, the data is visualized as a contact matrix.

### Region Capture Micro-C (RCMC) Protocol

Micro-C developed by Stanley Hsieh | Annotated by Claudia Cattoglio & Viraat Goel, & Capture-optimized by Viraat Goel

**I. Harvest, fix, and crosslink cells from cell culture[1,2].**

**1.** Culture cells in the appropriate conditions and medium.

**2.** PBS wash and trypsinize cells to harvest them. Count cells after inactivating the trypsin with media, and pellet cells by centrifugation for 5 min at 800xg at room temperature.

- Though not necessary, it is recommended to **perform a PBS wash of the cells post-centrifugation** to ensure all possible media and trypsin have been removed before proceeding to crosslinking.
- **Remove the DSG from the 4C fridge** during these steps, allowing it to equilibrate to room temp. Once you know the number of cells you will be harvesting (post-counting), you can simultaneously begin preparing your crosslinking solution for step 3.

*If simultaneously crosslinking with DSG and formaldehyde (if not, skip to the bottom of this page)*

**3.** Freshly prepare a DSG[3] (long crosslinker, 7.7 Å) 300-mM stock solution (100X) in DMSO (MW=326.26; 20 mg in 200 μL DMSO). Dilute 100 times with room-temperature PBS (200 μL to 19.8 mL).

- Each unused vial of DSG should contain 50 mg, which is sufficient for 50 million cells when dissolved in 500 uL of DMSO and then in 50 mL of PBS.

**4.** Resuspend cell pellet in the long crosslinker solution at a concentration of $1x10^6$ cells/mL. Incubate for 35 minutes at room temperature with mixing.

- Resuspend the pellet first with a 1-ml low-retention tip and then add the rest of the media with a larger pipette without touching the cells.
- Make sure that mixing is done gently – harsh nutation can cause cell damage and may result in significant cell loss.

**5.** Add 16% formaldehyde (dropwise, but still rapidly) to a final concentration of 1% (e.g. 2 mL for a 30 mL samples) and allow crosslinking to continue incubating for 10 min at room temperature with mixing.

- Add the formaldehyde in the fume hood; if you have several samples, staggering crosslinking may be best.

**6.** Add Tris pH 7.5 dropwise to a final concentration of 0.375 M to quench the reaction (e.g. for a 30 mL sample, 18 mL using 1 M Tris or 6.9 mL using 2 M Tris).
Incubate for 5 min at room temperature. Centrifuge for 5 min at 850xg at 4°C. Aspirate supernatant.

*If separately crosslinking with DSG and formaldehyde*

**3.** Prepare enough base media (w/o FBS) or PBS to resuspend cells at a concentration of $1x10^6$ cells/mL (max. 30 mL in 50 mL tube for adequate mixing) and add 16% formaldehyde to a final concentration of 1% (e.g. 2 mL for 30 mL sample) in the fume hood.

- Add the formaldehyde in the fume hood.

**4.** Resuspend cells in the fixation media. Incubate for 10 min at room temperature with mixing.

- Resuspend the pellet first with a 1-ml low-retention tip and then add the rest of the media with a larger pipette without touching the cells.
- Make sure that mixing is done gently – harsh nutation can cause cell damage and may result in significant cell loss.

**5.** Add Tris pH 7.5 dropwise to a final concentration of 0.375 M to quench the reaction (e.g. for a 30 mL sample, 18 mL using 1 M Tris or 6.9 mL using 2 M Tris).
Incubate for 5 min at room temperature. Centrifuge for 5 min at 850xg at 4°C. Aspirate supernatant.

**6.** Wash cells twice with 1X cold PBS at a concentration of $1x10^6$ cells/mL. Each time, centrifuge for 5 min at 850xg at 4°C and aspirate supernatant.

**7.** Freshly prepare a DSG[3] (long crosslinker, 7.7 Å) 300-mM stock solution (100X) in DMSO (MW=326.26; 20 mg in 200 μL DMSO). Dilute 100 times with room-temperature PBS (200 μL to 19.8 mL).

- Each unused vial of DSG should contain 50 mg, which is sufficient for 50 million cells when dissolved in 500 uL of DMSO and then in 50 mL of PBS.

**8.** Resuspend cell pellet in the long crosslinker solution at a concentration of $1x10^6$ cells/mL. Incubate for 45 minutes at room temperature with mixing.

- Resuspend the pellet first with a 1-ml low-retention tip and then add the rest of the media with a larger pipette without touching the cells.
- Make sure that mixing is done gently – harsh nutation can damage cells and may result in significant cell loss.

**9.** Add Tris pH 7.5 dropwise to a final concentration of 0.375 M to quench the reaction (e.g. for a 30 mL sample, 18 mL using 1 M Tris or 6.9 mL using 2 M Tris).
Incubate for 5 min at room temperature. Centrifuge for 5 min at 850xg at 4°C. Aspirate supernatant.

*Regardless of whether crosslinking was done simultaneously or separately, proceed below:*
**10.** Wash cells with 1X cold PBS at a concentration of $1x10^6$ cells/mL. Centrifuge for 5 min at 850xg at 4°C. Aspirate supernatant.

**11.** Resuspend cell pellet to a desired volume of cold PBS to aliquot the sample into multiple tubes (e.g. $30x10^6$ cells resuspended to 6 mL and aliquoted to 6 tubes, 1 ml each, will yield $5x10^6$ cells/tube). **Re-count cells** to gauge loss during crosslinking and determine the total number of feasible aliquots.

- Re-counting here is critical for establishing consistency in how many cells actually enter the rest of the protocol.

**12.** Aliquot the sample. Centrifuge for 5 min at 850xg at 4°C. Aspirate supernatant.
Snap freeze cell pellets in liquid nitrogen. Store @ -80°C until needed (up to several years).

#### II. Digest crosslinked chromatin with Micrococcal Nuclease (MNase)

**1.** Prepare fresh, "complete" MB#1:

| Stock Tot: | 5 mL | 10 mL | Final |
|---|---|---|---|
| MB#1 | 4850 μL | 9700 μL | 50 mM NaCl, 10 mM Tris, 5 mM $MgCl_2$, 1 mM $CaCl_2$, 0.2% NP-40, 1X PIC |
| 10% NP-40 | 100 μL | 200 μL | Use protein-grade NP-40 \| **make sure to use NP-40 Alternative, not NP-40** |
| 100X PIC | 50 μL | 100 μL | Protease inhibitor cocktail dissolved in MB#1, aliquoted, and stored in -20°C. |

**2.** Thaw a cell pellet of the desired size (1 or $5x10^6$ cells) in ice and resuspend in complete MB#1 at a concentration of $1x10^6$ cells/100 μL.
Incubate for 20 min on ice.
Centrifuge at ~1,750xg for 5 min at 4°C. Discard supernatant, leaving ~10-20 μL behind.

- Supernatant removal should be done with a P200 pipet tip; for larger volumes, append the tip to a P1000 tip.
- Spins can be performed at <=3,000xg until DNA extraction as long as a pellet clearly forms with little nuclear loss in the suspension. Centrifugation at <=2,000xg should be sufficient until at least proximity ligation.

**3.** Wash nuclei pellet in complete MB#1 at a concentration of $1x10^6$ cells/100 μL, resuspending the pellet up & down with a pipette. Centrifuge at ~1,750xg for 5 min at 4°C. Discard supernatant with a P200 pipette tip, removing as much liquid as possible without disturbing the pellet.

- Thaw MNase; partially used aliquots are in the -20C (use no more than 3 times), fresh aliquots are in the -80C

**4.** Resuspend nuclei pellet in complete MB#1 at a concentration of $1x10^6$ cells/100 μL, pipetting up & down.

**5.** Add the appropriate amount of MNase (as determined by an MNase titration experiment on the same sample batch) to digest chromatin to an 80-90% monomer / 20-10% dimer ratio.
Vortex briefly and incubate using the same conditions used in the MNase titration (usually 10-**20** min at 37°C with shaking at 850-1,000 RPM using a thermomixer).

- This step is critical, so double-check the chosen MNase amount. Underdigestion is bad, overdigestion is worse.

**6.** Transfer the nuclei back onto ice and promptly stop the reaction by adding 500 mM EGTA to a final concentration of 4 mM (0.8 μL for 100 μL). Vortex briefly to mix.
Incubate for 10 min at 65°C to ensure complete inactivation of the MNase.
Centrifuge at 1,750-2,000xg for 5 min at 4°C. Discard supernatant with a P200 tip.

- EGTA is a stronger $Ca^{2+}$ chelator than EDTA

**7.** Wash/rinse nuclei pellet in 1 mL of cold MB#2 twice.
Centrifuge for 5 min at 1,750-2,000xg at 4°C. Discard supernatant with a P200 pipette tip.
After the second wash, try and remove as much liquid as possible without disturbing the pellet.

#### III. Repair fragment ends

##### 1. End chewing

| Stock | Tot: | 5x10$^6$ cells 100 μL | 1x10$^6$ cells 50 μL | Final |
|---|---|---|---|---|
| Chromatin | | pellet | pellet | |
| $H_2O$ (first) | | 50 μL | 25 μL | |
| 10X NEBuffer 2.1 | | 10 μL | 5 μL | 50 mM NaCl, 10 mM Tris-HCl pH 7.5, 10 mM $MgCl_2$, 100 μg/mL BSA |
| 10 mM ATP | | 20 μL | 10 μL | 2 mM ATP |
| 100 mM DTT | | 5 μL | 2.5 μL | 5 mM DTT (freshly prepare 0.01543 g DTT in 1 mL HEPES buffer) |
| 10 U/μl T4 PNK (last) | | 5 μL | 2.5 μL | ~25-50 U / 300 pmol of DNA |

Can prepare a Master Mix of all reagents except the PNK

Incubate for 15 minutes at 37°C, shaking @ 1,000 RPM.

- T4 PNK catalyzes the transfer of a phosphate to the 5' ends of DNA and removes phosphates from 3' ends.

Add 5U/μL Klenow Fragment 10 μl 5 μL ~1 U /1μg of DNA

Incubate for 15 minutes at 37°C, shaking @ 1,000 RPM.

- In the absence of dNTPs, the Klenow fragment only possesses a 3'-5' exonuclease activity, creating a single-stranded DNA that can be labeled with biotin in the next step.

##### 2. End labeling

| Stock | Tot: | 5x10$^6$ cells 150 μL | 1x10$^6$ cells 75 μL | Final |
|---|---|---|---|---|
| Chromatin | | 100 μL | 50 μL | 33 mM NaCl, 1.3 mM ATP (from previous step) |
| 1 mM Biotin-dATP | | 10 μL | 5 μL | \|-> dNTPs: 66 μM each |
| 1 mM Biotin-dCTP | | 10 μL | 5 μL | \| |
| 10 mM dTTP | | 1 μL | 0.5 μL | \| |
| 10 mM dGTP | | 1 μL | 0.5 μL | \| |
| 10X T4 DNA ligase buffer | | 5 μL | 2.5 μL | 23 mM Tris-HCl pH 7.5, 10 mM $MgCl_2$, 0.33 mM ATP, 6.67 mM DTT |
| 20 mg/mL BSA (200X) | | 0.25 μL | 0.125 μL | 100 μg/mL BSA |
| $H_2O$ | | 22.75 μL | 11.375 μL | |

Can prepare a Master Mix of all of these reagents

Incubate for 45 minutes at 25°C with interval mixing @ 1,000 RPM.

- Use a thermomixer that allows interval mixing: 1 minute of shaking, followed by 3 minutes still, repeated.

Add 500 mM EDTA to a final concentration of 30 mM (9 μL in 150 μL) to quench the reaction.

Briefly vortex and incubate at 65°C for 20 minutes.

Centrifuge at 1,750-2,000xg for 5 minutes at 4°C. Discard supernatant with P200 tips.

Rinse once with 1 mL of cold MB#3, pipetting up and down.

Centrifuge at 1,750-2,000xg for 5 minutes at 4°C. Discard supernatant with P200 tips.

#### IV. Proximity ligation and purge of unligated ends

**1.** Proximity ligation

| Stock | Tot: | 5x$10^6$ cells **500 μL** | 1x$10^6$ cells **250 μL** | Final |
|---|---|---|---|---|
| Chromatin | | pellet | pellet | |
| $H_2O$ | | 422.5 μL | 211.25 μL | |
| 10X T4 DNA ligase buffer | | 50 μL | 25 μL | 50 mM Tris-HCl, 10 mM $MgCl_2$, 1 mM ATP, 10 mM DTT |
| 20 mg/mL BSA (200X) | | 2.5 μL | 1.25 μL | 100 μg/mL BSA |
| 400 U/μL T4 DNA ligase | | 25 μL | 12.5 μL | Add this last |

Incubate for at least 2.5 hours (overnight is alright) @ 25°C with slow rotation using a gentle nutator.

Centrifuge at 3,000xg for 5 minutes at 4°C. Discard supernatant with P200 tips.

**2.** Removal of biotin-dNTPs from unligated ends

| Stock | Tot: | 5x$10^6$ cells **200 μL** | 1x$10^6$ cells **100 μL** | Final |
|---|---|---|---|---|
| Chromatin | | pellet | pellet | |
| $H_2O$ | | 170 μL | 85 μL | |
| 10X NEBuffer#1 | | 20 μL | 10 μL | 10 mM Bis-Tris-Propane-HCl, 10 mM $MgCl_2$, 1mM DTT |
| 100 U/μL Exonuclease III | | 10 μL | 5 μL | |

Incubate at 37°C for 15 minutes with interval mixing (1 minute shaking @ 1,000 RPM, 3 minutes still).

**3.** Reverse crosslinking

| Stock | Tot: | 5x$10^6$ cells **265 μL** | 1x$10^6$ cells **133 μL** | Final |
|---|---|---|---|---|
| Chromatin | | 200 μL | 100 μL | |
| 20 mg/ml Proteinase K | | 26 μL | 13 μL | 2 mg/mL |
| 10% SDS | | 26 μL | 13 μL | 1X |
| 5 M NaCl | | 10.4 μL | 5.2 μL | 250 mM |
| 10 mg/mL RNaseA | | 2.6 μL | 1.3 μL | 0.1 mg/mL |

*Notes:* Don't make a master mix since the SDS would be extra concentrated. Use filter tips once RNaseA has been added.

Incubate @ 65°C overnight.

- If incubated in a water bath, the sample lids should be carefully covered with Parafilm to avoid any lid popping open accidents. Beware that this can cause ink smudging, so label carefully and clearly to distinguish samples.

#### V. Di-nucleosomal DNA purification

**1.** Phenol-Chloroform:Isoamylic alcohol (PCI) extraction *(perform all steps in the fume hood)*

i. Transfer sample to labeled phase-lock tubes.

- A quick spin may be necessary pre-transfer to remove gel from the tube lid.

ii. Add an equivalent volume of PCI to the sample volume (if using 5x$10^6$ cells, add 265 μL of PCI to 265 μL of sample).

- PCI often comes with a top "containment" layer of liquid – avoid pipetting from this top layer and instead pipette from the PCI solution beneath it.

iii. Vortex for 20 seconds and then spin for 15 minutes at 19,800xg at room temperature.

iv. Transfer the upper layer to a new low-retention tube.

v. *(Optional)* Add 50 μL TE to each phase-lock tube and vortex again. Spin for 10 minutes at 19,800xg at room temp.

- This second spin will help with the removal of any residual aqueous layer DNA.

vi. Transfer the upper layer to the previously used low-retention tube.

**2.** DNA purification by ethanol precipitation (optional, usually skipped by Viraat)

i. Add 0.1X volumes of 3M sodium acetate and 2.5X volumes of cold 100% ethanol to the sample volume.

ii. Invert the tube and incubate for at least 1 hour at -80°C (overnight is alright but not necessary).

iii. Spin at 19,800xg for 15 minutes @ 4°C.

iv. Discard ethanol with a P200 pipette and wash the pellet with 1 mL of cold 80% ethanol.

- A pellet may be faint or impossible to see – that's okay! This is likely the result of not starting with many cells.

v. Spin at 19,800xg for 5 minutes @ 4°C.

vi. Remove as much ethanol as possible (if needed spin the tube briefly once again) and air dry the pellet for 4 minutes @ 37°C with an open lid.

- The pellet shouldn't have a discernable layer of ethanol on it, but it also shouldn't be super dry.

vii. Add 50 μL of TE buffer on top of the pellet and let it dissolve @ 37°C for 30 minutes. Intermittently vortex and tabletop spin down to help dissolve all DNA.

*Cast a gel (usually 100 mL of a 1% gel made using regular agarose with SYBR Safe stain in the 12-well turquoise cassettes) at this time so it can set as you use the DCC kit.

**3.** ZymoClean: DNA purification with the Zymo DNA Clean & Concentrator (e.g. DCC-25) kits

Follow kit instructions with the minor alterations noted below.

- Be mindful of the maximum DNA binding capacity of the column you are using, and save / test flow-through in case you are unsure

i. Add 300 μL of Zymo DNA binding buffer per 50 μL of DNA sample and load to the column.

ii. Wash twice with 400 μL of wash buffer.

- Perform a quick "dry spin" after the second wash to remove any residual wash buffer.

iii. Elute twice, each time with 25 μL of EB (50 μL total).

- To maximize yield, warm up elution buffer to 60-70 °C before adding it to the middle of the column filter
- Be mindful of the recommended elution volume for the column, and adjust / increase volumes accordingly to ensure saturation.

**4.** Size-selection of di-nucleosomal DNA

i. Quantify the sample concentration using a Nanodrop or Qubit

- This should help you benchmark extraction yield and can help avoid "overloading" a gel well. For example, loading 1-5 ug of DNA per well of a standard 10-well gel should be alright, but 7+ ug may lead to gel running issues.

ii. Add 10X Orange G loading dye to the sample in greater than 10X conc. (e.g. add 12 μL dye to 50 μL sample).

- This is necessary to densify the sample so it sits in the gel well and doesn't float out.
- Do **not** vortex samples to mix; this can trap air bubbles in your sample and cause it to float out of the well

iii. Load the sample(s) onto the gel; if necessary, load to two large wells. Also load 1kb+ DNA ladder.

iv. Run the gel at 120V for 60 minutes.

v. Use the GelDoc Imager to image the gel and cut out the dinucleosomal DNA band,

- Cut the band between 250-400 bp or 200-350 bp. Avoid cutting below 200 bp to eliminate monomers.

***Note:*** A sample gel and extraction from Berkeley is shown below. The cut strayed a bit below 200 bp and should have been made a little higher.

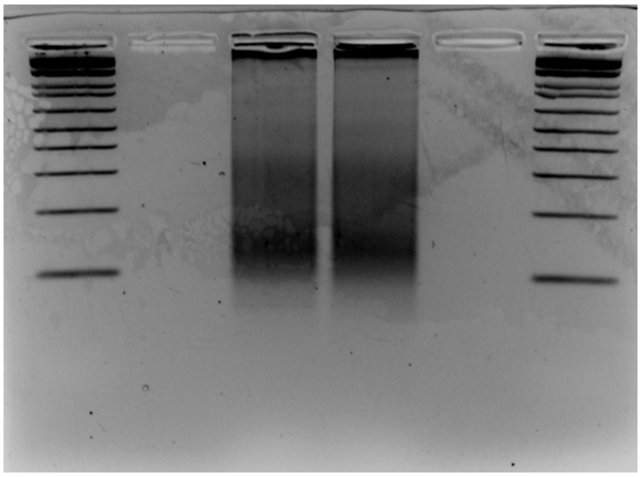

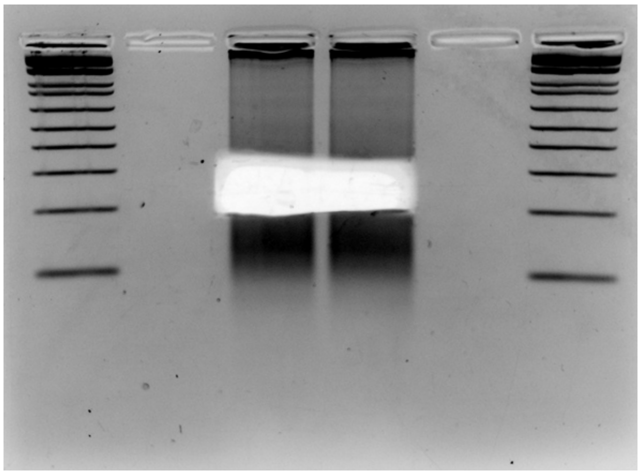

**5.** Gel purification with the Zymo gel extraction kit

Follow the kit instructions but wash twice with 400 μL of wash buffer.

- Perform a quick "dry spin" after the second wash to remove any residual wash buffer.

Elute twice, each time with 9 μL of EB (18 μL total).

- To maximize yield, warm up elution buffer to 60-70 °C before adding it to the middle of the column filter.

***Note:*** When performing Micro-C for the first time, quantify with Qubit at this point. The DNA amount should be at least 10 ng for a good result.

#### VI. Library preparation

**1.** End polishing[4] (assemble the reaction at room temperature)

| Stock | Tot: 25 μL | |
|---|---|---|
| Input DNA | 17 μL | |
| 10X End-it buffer | 2.5 μL | |
| 10X ATP | 2.5 μL | |
| 10X dNTPs | 2.5 μL | |
| End-it enzyme mix | 0.5 μL | T4 polymerase + Klenow |

Incubate for 45 minutes @ 25°C (without mixing).
Inactivate the enzyme mix by incubating for 10 minutes @ 65°C.

*– This is a good point to stop. Store your sample at 4 ℃ or -20 ℃ to proceed the day after. –*

**2.** Streptavidin purification

*Beads pre-wash:*

i. For x number of samples, add 1 mL of 1X TBW[5] to 2.5/5 μL (for 1/5x$10^6$ cells, respectively) of C1 or T1 streptavidin beads **into x separate tubes** (the sample(s) and the beads are combined after a couple wash steps, not right now).
   - Also, using more beads for pulldown can improve yields (test this), likely because 2.5/5 μL of beads may get saturated (there are multiple biotin per Micro-C DNA fragment). We currently use 25-30 μL of beads per 5M cells of input for a sample.

ii. Nutate for 2 minutes, followed by the magnetic stand for 30-60 seconds.

iii. Carefully remove supernatant with a P200 tip attached to a P1000 tip without disturbing the beads.

iv. Resuspend the beads in 150 µL of 2X BW.

v. Bring the volume of the samples to 150 µL with $ddH_2O$ (25 µL of extracted / purified DNA + 125 µL of water).

vi. Add the beads to the sample(s). Mix the DNA and the beads for 20+ minutes at room temperature (O/N mixing also works well).

*Sample wash steps:*

vii. Spin briefly in a benchtop microcentrifuge and add 950 µL of 1X TBW.
viii. Invert the tube and shake @ RT, 1200 RPM for 5 minutes on a mixer.
ix. Spin briefly in a bench microfuge and place on the magnet for 30-60 seconds.
x. Carefully remove the supernatant and add 950 µL of 1X TBW to repeat the wash a second time.
xi. Spin briefly in a benchtop microcentrifuge, place on the magnet for 30-60 seconds and carefully remove the supernatant.
xii. With the tube on the magnetic stand, slowly rinse the beads with 10 mM Tris-HCl pH 7.5 without disturbing the beads.
xiii. Remove 10 mM Tris once the End-Repair mix from the next step is ready
- Never let your beads dry out![6]

**3.** End Repair and A-tailing

| **Stock** | **Tot: 30 µL** |
|---|---|
| $H_2O$ | 25 µL |
| End Prep Reaction Buffer | 3.5 µL |
| End Prep Enzyme Mix | 1.5 µL |

***Note:*** Steps 3 & 4 to the left have halved volumes compared to the **NEBNext kit protocol**. If you're combining samples to have >5M cells of material per tube, **consider using doubled volumes** for these steps; the only exception to doubling is the amount of adapter (NEBNext calls for 2.5 µL of adapter).

i. Remove the 10 mM Tris from the beads above and resuspend them in the End Repair mix by either pipetting or vortexing.
- Make sure to avoid bubbles.

ii. Incubate for 30 minutes @ 20°C with interval mixing at 1,000 RPM (1 minute on every 3 minutes off).

iii. Inactivate the enzymes by incubating for 30 minutes @ 65°C.

iv. Transfer to ice.

**4.** Adapter ligation[7]

| **Stock** | **Tot: 46 µL** |
|---|---|
| Input DNA | 30 µL |
| NEB Illumina adapter (first)[8] | 0.5 µL |
| Ligation Master Mix | 15 µL |
| Ligation enhancer (PEG 8000) | 0.5 µL |

***Note:*** Dilute the adapter in $ddH_2O$ or Tris pH 7.5-8.0 based on how much DNA you have. Look @ NEB's recommendations for the NEBNext prep.
Also, the Ligation Master Mix and Enhancer can be mixed prior to addition to samples. The L.M.M. is especially viscous, so mix everything thoroughly.

Vortex briefly and incubate for 30 minutes @ 20°C with interval mixing at 1,000 RPM (as before).

Add USER enzyme 1.5 µL

Incubate for 15 minutes @ 37°C with interval mixing at 1,000 RPM (as before).

**5.** Bead wash

i. Add 950 µL of 1X TBW, invert the tube, and mix at 1,200 RPM at @ RT for 3 minutes on a mixer.

ii. Spin the tube briefly, transfer to the magnetic stand, and carefully remove the supernatant.

iii. Repeat the 1X TBW wash (steps i and ii) a second time.

iv. Slowly rinse the beads with 10 mM Tris-HCl pH 7.5 without disturbing the beads (keep in magnetic stand).

v. Resuspend the beads in at least 20 µL of EB buffer (e.g. from the Zymo kit); you can do more, e.g. 50 µL or up to 200-300 µL depending on desired # of test PCRs and pool PCRs. Briefly vortex and spin to gather them.

*– This is a good point to stop if needed. Store your sample at 4℃ or -20℃ to proceed the day after. –*

**6.** Test PCR run[9]. Assemble the reaction in 200-μL PCR tubes in ice (10 μL final reaction volume).

| **Stock** | **Tot:** | **10 μL** |
|---|---|---|
| Streptavidin beads (add last) | | 1 μL |
| $H_2O$ | | 3.5 μL |
| 2X Q5 NEB mix | | 5 μL |
| 10 μM universal primer[10] | | 0.25 μL |
| 10 uM index primer[10] | | 0.25 μL |

***Note:*** NEBNext recommends **4x** these primer volumes; if following NEBNext's guidance, add 1 μL of each primer and only 2 μL of water.

Mix the reaction by either pipetting up and down or vortexing.

| PCR conditions: | Q5 NEB | KAPA HiFi |
|---|---|---|
| Start with: | 98°C, 30" | 98°C, 45" |
| 11-15 cycles of: | 98°C, 10" | 98°C, 15" |
| | 65°C, 1'15" | 60°C, 30" |
| | | 72°C, 30" |
| Ending with: | 65°C, 5' | 72°C, 60" |
| | 4°C on hold | 4°C on hold |

***Note:*** Can alternatively use other polymerases (e.g. KAPA HiFi Hotstart ReadyMix; if so, change the thermocycler program accordingly. Both the Q5 & KAPA programs are shown at the left

Add 3 μL of 10X Orange-G and load all of the sample on a regular 1-1.5% agarose gel side-by-side with 1Kb-plus DNA ladder.

- No need to remove beads, they will stay in the well without disturbing DNA migration.

Run the gel at 120 V for about 40 minutes.

Take a picture and calculate how much material you obtained by amplification using the ladder for comparison

- For example, let's say we have a band estimated at 10 ng total. We will need 50 ng for regular Micro-C sequencing (or 500 ng for target enrichment) – 5 times more – but we only used a 20th of the beads for this test run. So, we can "afford" 4-fold less yield, corresponding to 2 less PCR cycles. Perform 9 instead of 11 cycles of amplification for the whole sample.
- Also use this gel to check for the presence of adapter dimers (increase with more test PCR cycles) and primer dimers (decrease with more test PCR cycles).

**7.** Pool PCR run. Assemble the reaction in 200-μL PCR tubes in ice (50 μL total volume).

| **Stock** | **Tot:** | **50 μL** |
|---|---|---|
| Streptavidin beads (add last) | | 19 μL |
| $H_2O$ | | 3.5 μL |
| 2X Q5 NEB mix | | 25 μL |
| 10 μM universal primer[10] | | 1.25 μL |
| 10 uM index primer[10] | | 1.25 μL |

***Note:*** NEBNext recommends **4x** these primer volumes; if following this, add 5 μL of each primer and 29 μL of 2X enzyme for a 58 μL reaction, or split across multiple 50 μL reactions.

***Note:*** Can alternatively use other polymerases (e.g. KAPA HiFi Hotstart ReadyMix; if so, change thermocycler program accordingly.

Mix the reaction by either pipetting up and down or vortexing.

| PCR conditions (as before, Q5 shown) | |
|---|---|
| Start with: | 98°C, 30" |
| 6-9 cycles[11] of: | 98°C, 10" |
| | 65°C, 1'15" |
| Ending with: | 65°C, 5' |
| | 4°C on hold |

***Note:*** Although the C1 & T1 bead datasheets note that beads with similar chemistries (namely, M280 beads with the same core chemistry as T1 beads) can significantly inhibit PCRs when >75 μg of beads are present in a single reaction, I have successfully tested 30 μL (300 μg) of both C1 and T1 beads in individual PCR reactions and found no noticeable inhibition. It should also be noted that T1 beads outperformed the C1 beads by 1.5-3-fold (about 1-2 cycles' worth of amplification).

Post-PCR:

1. Transfer all of the PCR reaction to a low-retention 1.5 mL tube.
2. Place the tube on a magnetic rack and allow the streptavidin beads to move to the side of the tube until the liquid is clear.
3. Move the liquid to a new low-retention 1.5 mL tube and proceed.

> ***Note:*** You could proceed and mix the AmPure and streptavidin beads together; however, for the sake of bead separation, this allows the removal of streptavidin beads before adding the cleanup beads.

**8.** 0.9X Ampure XP beads purification.[12]

I. Allow the Ampure XP beads to equilibrate to RT and mix them well.
II. For each 50 μL of post-PCR sample, add 45 μL of beads and pipet up and down until thoroughly resuspended.
III. Incubate at RT for 15 min to allow the DNA to bind to beads.
IV. Place the tube onto the magnetic rack and incubate until solution is clear.
V. Remove and discard the supernatant without disturbing the beads.
VI. Add 200 μL of freshly prepared 80% EtOH without disturbing the beads.
VII. Incubate the solution for 30 seconds while still on the magnetic rack.
VIII. Remove and discard the EtOH without disturbing the beads.
IX. Repeat the EtOH wash steps (VI-VIII).
X. Remove residual ETOH using a P20 and/or vacuum without disturbing the beads.
XI. Air-dry the beads for 5 minutes (do not overdry the beads!).
XII. Remove the tube from the magnetic rack & add at least 25 μL of elution buffer or water to the sample.
    a. The aim is to have a final concentration of DNA between 4 and 20 nM (in theory, you should use 50 μL for 1000 fmol to get 20 nM, but we account for 50% of material loss).
XIII. Thoroughly re-suspend the beads by pipetting up and down 10-20x.
XIV. Incubate at RT for 2 minutes to allow DNA to elute off of the beads.
XV. Place the tube back on the magnetic rack and incubate until the solution is clear.
XVI. Transfer the supernatant to new low-bind tube.

> ***Note:*** If NOT proceeding to Capture, it may be worthwhile to do **2** consecutive rounds of AmPure cleanup to ensure contaminating adapter dimers are eliminated.

**9.** Qubit and qPCR quantify samples.

- Also, determine the library fragment distribution using a Fragment Analyzer.

**10.** Pool barcoded samples (and submit for sequencing if doing regular Micro-C, or to target enrichment if doing Tiled-Micro-C).

- Barcoded samples are regularly pooled at a 1:1 molar ratio.
- If doing regular Micro-C:
    - Consider using ~ 1 pmol total DNA, splitting this amount between barcoded samples (e.g., 5 samples, 100 fmol each; 20 samples, 50 fmol each).
    - Determine the sequencing technology you would like to use and its associated requirements (sample volume & concentration). Submit your pool and enjoy the resulting data!
        - Paired-end sequencing with 35-75 bp read lengths is recommended.
- If doing Tiled-Micro-C, segue into Twist's Target Enrichment Protocol and follow it verbatim
    - **Exception #1**: Add a test PCR to confirm how many amplification cycles are necessary.
        - Often, fewer cycles than recommended are necessary. E.g. Capturing 3 Mb total from a 2-4 μg Capture input should meet sequencing submission needs in 6-7 cycles.
    - **Exception #2**: If working with mouse samples, replace Twist's Blocker Solution (which is human-specific) with Mouse Cot-1 DNA to similarly prevent nonspecific binding to probes.
    - **Note:** Multiple Capture panels can be combined (e.g. 3 1-Mb panels) – just combine them, vacuum dry them, and resuspend them in the protocol-specified volume before proceeding.

**Materials and Reagents**

| Item | Manufacturer / Supplier | Catalog # |
|---|---|---|
| Formaldehyde | ThermoFisher | 28906 |
| DSG (disuccinimidyl glutarate) | ThermoFisher | 20593 |
| Tris buffer | K-D Medical | RGE-3370 |
| Low-retention pipet tips | VWR | 76322 |
| NP-40 Alternative | Millipore Sigma | 492018 |
| Protease Inhibitor Cocktail (EDTA-free) | Sigma-Aldrich | 5056489001 |
| MNase (micrococcal nuclease) | Worthington Biochem | LS004798 |
| EGTA | bioWORLD | 40520008 |
| BSA (bovine serum albumin) | Sigma-Aldrich | B8667 |
| T4 PNK (polynucleotide kinase) | New England BioLabs | M0201 |
| ATP | ThermoFisher | R1441 |
| DTT (dithiothreitol) | Sigma-Aldrich | 10197777001 |
| DNA Polymerase I Klenow Fragment | New England BioLabs | M0210 |
| dTTP | Jena Bioscience | NU-1004 |
| dGTP | Jena Bioscience | NU-1003 |
| Biotin-14-dATP | Jena Bioscience | NU-835-BIO14 |
| Biotin-11-dCTP | Jena Bioscience | NU-809-BIOX |
| EDTA | Invitrogen | 15575020 |
| T4 DNA Ligase | New England BioLabs | M0202 |
| Exonuclease III | New England BioLabs | M0206 |
| Proteinase K | Viagen Biotech | 501-PK |
| RNase A, DNase- and proteinase-free | ThermoFisher | EN0531 |
| SDS (sodium dodecyl sulfate) | Sigma-Aldrich | L3771 |
| Tris-EDTA buffer | Sigma-Aldrich | 93283 |
| Phenol:Chloroform:Isoamylic alcohol (PCI) | Sigma-Aldrich | P2069 |
| 5PRIME Phase Lock Gel Light tubes | Quantabio | 2302820 |
| Low-retention tubes (DNA LoBind) | Eppendorf | 0030108418 |
| DNA Clean & Concentrator kit | Zymo Research | D4034 |
| Agarose | VWR | 97062 |
| Gel Purification kit | Zymo Research | D4008 |
| Qubit dsDNA HS Assay kit | Invitrogen | Q33231 |
| End-it DNA End-Repair kit | Lucigene | ER81050 |
| Dynabeads MyOne Streptavidin C1 | Invitrogen | 65001 |
| Dynabeads MyOne Streptavidin T1 | Invitrogen | 65601 |
| Tween-20 | Sigma-Aldrich | P8074 |
| NEBNext Ultra II kit | New England BioLabs | E7645 |
| Multiplex Oligos for Illumina Primer Set 1 | New England BioLabs | E7335 |
| KAPA HiFi HotStart ReadyMix | Roche | 07958927001 |
| Q5 High-Fidelity DNA Polymerase | New England BioLabs | M0491 |
| AmPure XP beads | Beckman Coulter | A63880 |
| Capture Custom Panels | Twist Bioscience | 101001 |
| Standard Hybridization Mix | Twist Bioscience | 104178 |
| Universal Blockers | Twist Bioscience | 100578 |
| Mouse Cot-1 DNA (only if using mouse samples) | Invitrogen | 18440016 |
| Streptavidin Binding Beads | Twist Bioscience | 100983 |
| Equinox Library Amplification Mix | Twist Bioscience | 104178 |
| DNA Purification Beads | Twist Bioscience | 100983 |

#### Stock Solutions

**100X cOmplete protease inhibitor cocktail (PIC) (EDTA-free)**
Dissolve one EDTA-free PIC tablet in 500 µL of MB#1 buffer, separate into 50-100 µL aliquots, and store at -20°C.

**10 mM Tris-HCl pH 7.5 (100 mL; elution buffer)**
1 mL of 1 M Tris-HCl pH 7.5 stock to 99 mL of $ddH_2O$. Sterile filter.

**1X TE (Tris-EDTA)**
1 mL of 10X TE (100 mM Tris-HCl pH 8.0 and 10 mM EDTA) stock to 9 mL of $ddH_2O$. Sterile filter. The final buffer contains 10 mM Tris-HCl pH 8.0 and 1 mM EDTA. Can also purchase a prepared 1X solution.

**20 U/µL MNase**
Dissolve MNase to 20 U/µL in 10 mM Tris-HCl pH 7.5.
Aliquot into 20 µL aliquots in tubes and store at – 80°C. Upon first use, re-freeze at – 20°C and use another couple of times (maximum 3 times per aliquot).

**MB#1**

| Stock | Final | For 500 mL |
|---|---|---|
| 5 M NaCl | 50 mM | 5 mL |
| 1 M Tris-HCl pH 7.5 | 10 mM | 5 mL |
| 1 M $MgCl_2$ | 5 mM | 2.5 mL |
| 2.5 M $CaCl_2$ | 1 mM | 200 µL |

Bring volume to 500 mL with $ddH_2O$ and sterile filter.
Add NP-40 and protease inhibitors right before use. $CaCl_2$ will activate MNase right away.

**MB#2**

| Stock | Final | For 500 mL |
|---|---|---|
| 5 M NaCl | 50 mM | 5 mL |
| 1 M Tris-HCl pH 7.5 | 10 mM | 5 mL |
| 1 M $MgCl_2$ | 10 mM | 5 mL |

Bring volume to 500 mL with $ddH_2O$ and sterile filter.

***Note:*** Add BSA to a final concentration of 100 µg/mL to MB #2 & #3 to help with cell pelleting.

**MB#3**

| Stock | Final | For 500 mL |
|---|---|---|
| 1 M Tris-HCl pH 7.5 | 50 mM | 25 mL |
| 1 M $MgCl_2$ | 10 mM | 5 mL |

Bring volume to 500 mL with $ddH_2O$ and sterile filter.

**Orange G 6X gel loading dye**

| Stock | Final | For 50 mL |
|---|---|---|
| Orange G | 0.1% w/v | 500 mg |
| 1 M Tris-HCl pH 7.5 | 10 mM | 500 µL |
| 0.5 M EDTA pH 8.0 | 60 mM | 6 mL |
| Glycerol | 60% | 30 mL |

Bring volume to 50 mL with $ddH_2O$ and sterile filter. Orange G will migrate @ ~50 bp in a 1% agarose gel.

**Reverse crosslinking solution** (if resuspending cell pellets)

| Stock | Final | For 0.9 mL (6X) |
| --- | --- | --- |
| 1X TE | | 720 µL |
| 20 mg/ml Proteinase K, 20X | 1X | 45 µL |
| 10% SDS | 1% | 90 µL |
| 10 mg/ml RNaseA (100X) | 1X | 9 µL |
| 5 M NaCl | 1X | 36 µL |

**3M Sodium acetate pH 5.2 (500 mL)**

Dissolve 123.05 g of sodium acetate in 250 mL of dd$H_2O$. Adjust the pH to 5.2 with glacial acetic acid. Allow the solution to cool overnight. Adjust the pH once more to 5.2 with glacial acetic acid. Adjust the final volume to 500 mL with dd$H_2O$ and filter-sterilize.

**2XBW**

| Stock | Final | For 500 mL |
| --- | --- | --- |
| 5 M NaCl | 2 M | 200 mL |
| 1 M Tris-HCl pH 7.5 | 10 mM | 5 mL |
| 0.5 M EDTA pH 8.0 | 1 mM | 1 mL |

Bring volume to 500 mL with dd$H_2O$ and sterile filter.

**TBW (100 mL)**

| Stock | Final | For 250 mL |
| --- | --- | --- |
| 2XBW | 1X | 125 mL |
| 100% Tween-20 | 0.1% | 250 µL |

Bring volume to 250 mL with dd$H_2O$ and sterile filter.

**Notes**

1. Every batch of cross-linked cells needs to be tested for MNase digestion. Starting with a P10 will give you enough cells to run an MNase titration and also prepare a library
2. Use low-retention tubes and pipet tips every time you are pipetting nuclei/sample
3. EGS can also be used (long crosslinker, 16.1 Å, MW=326.26). In yeast, there is no difference between EGS and DSG, while Stanley has never tested EGS in mammalian cells.
4. It is useful to remove the single-stranded DNA that is created by the exonuclease before binding the ligation product to streptavidin, which could be sticky and non-specifically binding to them
5. The streptavidin beads have a really high capacity, so 5 μL will be more than enough when starting from $5x10^6$ cells.
6. The DNA is never eluted from the beads, and beads are carried over all the way through the PCR reaction.
7. NEB universal adapters have a U-shaped structure that impedes the formation of adapter dimers. Once ligated, the U-shape is removed by the USER enzyme, leaving the ends available for PCR primer annealing. The barcodes will be introduced using a specific PCR primer, so that you have the flexibility of re-amplifying your product with different barcodes, if needed.
8. Check NEB tables if you have low input amounts; you will need to dilute adapters.
9. You will first run an 11-cycle test PCR with only 1 μL of the sample to check the right size and estimate the PCR yield on agarose gel. Based on the yield, you will then perform a real PCR with hopefully a lower amount of cycles (typically ~8).
10. Official Illumina name. One of the primers is a universal one, while the other contains a unique barcode. From the NEB manual (available at: https://www.neb.com/-/media/nebus/files/manuals/manuale7335.pdf):

    *If fewer than 7 indexes are used in a lane for sequencing, it is recommended to use the following:*
    *Pool of 2 samples: Index #6 and 12*
    *Pool of 3 samples: Index #4, 6 and 12*
    *Pool of 6 samples: Index #2, 4, 5, 6, 7 and 12*
11. A pretty typical number of PCR cycles is 8, but this will need to be adjusted on the basis of the test PCR run.
12. Bring the AmPure beads to room temperature before use.

### Micrococcal Nuclease (MNase) Titration Protocol – For RCMC

Micro-C developed by Stanley Hsieh | Annotated by Claudia Cattoglio & Viraat Goel

*Notes:*

1. This titration protocol is a truncated version of the full **Region Capture Micro-C Protocol**.
2. Use low-retention tubes and tips every time you are pipetting the cell/nuclear pellets.

#### I. Harvest, fix, and crosslink cells from cell culture.

*Completed as per the Region Capture Micro-C protocol.*

#### II. Digest crosslinked chromatin with Micrococcal Nuclease (MNase)

**1.** Prepare fresh, "complete" MB#1:

| Stock Tot: | 5 mL | 10 mL | Final |
|---|---|---|---|
| MB#1 | 4850 µL | 9700 µL | 50 mM NaCl, 10 mM Tris, 5 mM $MgCl_2$, 1 mM $CaCl_2$, 0.2% NP-40, 1X PIC |
| 10% NP-40 | 100 µL | 200 µL | Use protein-grade NP-40 \| **make sure to use NP-40 Alternative, not NP-40** |
| 100X PIC | 50 µL | 100 µL | Protease inhibitor cocktail dissolved in MB#1, aliquoted, and stored in -20C. |

**2.** Thaw a cell pellet of the desired size (1 or 5x$10^6$ cells) in ice and resuspend in complete MB#1 at a concentration of 1x$10^6$ cells/100 µL.
Aliquot 100 µL of cells to 5x low-retention 1.5 mL Eppendorf tubes (if performing the titration on 1M cells)
Incubate for 20 min on ice.
Centrifuge at ~1,750xg for 5 min at 4°C. Discard supernatant, leaving ~10-20 µL behind.

- Supernatant removal should be done with a P200 pipet tip; for larger volumes, append the tip to a P1000 tip.

**3.** Wash nuclei pellet in complete MB#1 at a concentration of 1x$10^6$ cells/100 µL, resuspending the pellet up & down with a pipette. Centrifuge at ~1,750xg for 5 min at 4°C. Discard supernatant with a P200 pipette tip, removing as much liquid as possible without disturbing the pellet.

- **Thaw MNase**; partially used aliquots are in the -20C (use no more than 3 times), fresh aliquots are in the -80C
- If using 1M cell aliquots or if pipetting larger volumes is preferred, prepare a 1:20 dilution in 10 mM Tris-HCl, pH 7.5 (2 µL MNase + 38 µL buffer).

**4.** Resuspend nuclei pellet in complete MB#1 at a concentration of 1x$10^6$ cells/100 µL, pipetting up & down.

**5.** Add increasing amounts of MNase as follows:

**If digesting 1M cell aliquots**

**2 U:** 2 µL of a 1U/µL MNase dilution
**4 U:** 4 µL of a 1U/µL MNase dilution
**6 U:** 6 µL of a 1U/µL MNase dilution
**9 U:** 9 µL of a 1U/µL MNase dilution
**14 U:** 14 µL of a 1U/µL MNase dilution

**If digesting 5M cell aliquots**

**10 U:** 0.5 µL of the 20U/µL MNase stock
**20 U:** 1 µL of the 20U/µL MNase stock
**30 U:** 1.5 µL of the 20U/µL MNase stock
**45 U:** 2.25 µL of the 20U/µL MNase stock
**70 U:** 3.5 µL of the 20U/µL MNase stock

The target is to digest chromatin to an 80-90% monomer / 20-10% dimer ratio.
Briefly vortex the tubes and incubate for **20 min at 37°C** with shaking at 1000 rpm.
Prepare a 1.5% regular agarose gel.

**6.** Transfer the nuclei back onto ice and promptly stop the reaction by adding 500 mM EGTA to a final concentration of 4 mM (0.8 µL for 100 µL). Vortex briefly to mix.
Incubate for 10 min at 65°C to ensure complete inactivation of the MNase.
Centrifuge at 1,750-2,000xg for 5 min at 4°C. Discard supernatant with a P200 tip.

#### III. Isolate DNA and Visualize the Titration

**7.** Resuspend each pellet in 150 µL of Reverse Crosslinking solution, resuspending up and down.
Reverse cross-links for a minimum of 2 hours to overnight at 65°C.

**8.** Phenol-Chloroform:Isoamylic alcohol (PCI) extraction *(perform all steps in the fume hood)*

i. Bring sample volumes to 200 µL (add 50 µL) with 1X TE, and transfer sample to labeled phase-lock tubes.

- A quick spin may be necessary pre-transfer to remove gel from the tube lid.

ii. Add 200 µL of PCI to the sample volume.

- PCI often comes with a top "containment" layer of liquid – avoid pipetting from this top layer and instead pipette from the PCI solution beneath it.

iii. Vortex for 20 seconds and then spin for 15 minutes at 19,800xg at room temperature.

iv. Transfer the upper layer (~150 µL) to a new low-retention tube.

**9.** ZymoClean: DNA purification with the Zymo DNA Clean & Concentrator (e.g. DCC-25) kits

i. Add 5 volumes (1 mL) of DNA binding buffer and load to the column (you may have to load the column multiple times to load all of your sample).

ii. Wash twice with 400 µL of wash buffer.

- Perform a quick "dry spin" after the second wash to remove any residual wash buffer.

iii. Elute with 25 µL of elution buffer (EB).

**10.** Quantify the DNA by Nanodrop and check the size on a 1.5% regular agarose gel.

- This should help you benchmark extraction yield and can help avoid "overloading" a gel well. For example, loading 1-5 ug of DNA per well of a standard 10-well gel should be good, but 7+ ug may lead to gel running issues.

Add 10X loading dye to the samples. Using Orange G as a dye will minimally interfere with the size of the nucleosomal fragments.

Load the samples onto the gel; also load 1kb+ DNA ladder.

Run the gel (e.g. 120V for 60 min) and visualize it using a gel imager. **Pick the sample with the ideally-digested ratio of monomers to dimers** (a faint trimer band should be visible, while a tetramer band should not be visible).

**<u>Example Gel</u>**

This is a titration of mESC cells; a 5M cell pellet was **split into 1M cell aliquots** for this titration.

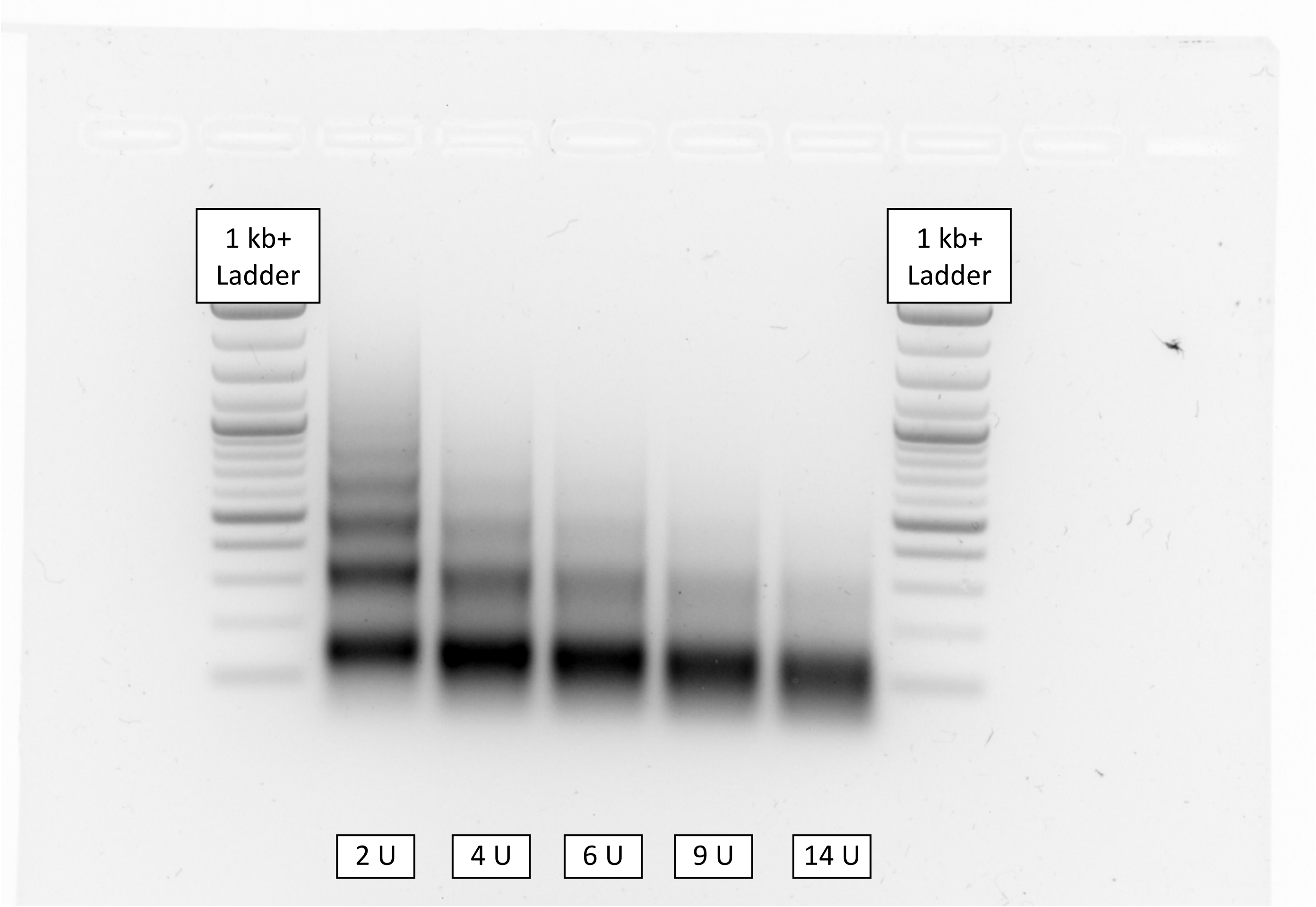


**In this case, the ideal MNase amount is:**

- Between <u>6-9 U for 1M cell samples</u>
- Between <u>30-45 U for 5M cell samples</u> (the 1M cell numbers here simply scaled up 5X)

#### Stock Solutions

**100X c*O*mplete protease inhibitor cocktail (PIC) (EDTA-free)**
Dissolve one EDTA-free PIC tablet in 500 µL of MB#1 buffer, separate into 50-100 µL aliquots, and store at -20°C.

**10 mM Tris-HCl pH 7.5 (100 mL)**
1 mL of 1 M Tris-HCl pH 7.5 stock to 99 mL of dd$H_2O$. Sterile filter.

**1X TE (Tris-EDTA)**
1 mL of 10X TE (100 mM Tris-HCl pH 8.0 and 10 mM EDTA) stock to 9 mL of dd$H_2O$. Sterile filter. The final buffer contains 10 mM Tris-HCl pH 8.0 and 1 mM EDTA. Can also purchase a prepared 1X solution.

**20 U/µL MNase**
Dissolve MNase to 20 U/µL in 10 mM Tris-HCl pH 7.5.
Aliquot into 20 µL aliquots in tubes and store at – 80°C. Upon first use, re-freeze at – 20°C and use another couple of times (maximum 3 times per aliquot).

**MB#1**

| Stock | Final | For 500 mL |
|---|---|---|
| 5 M NaCl | 50 mM | 5 mL |
| 1 M Tris-HCl pH 7.5 | 10 mM | 5 mL |
| 1 M $MgCl_2$ | 5 mM | 2.5 mL |
| 2.5 M $CaCl_2$ | 1 mM | 200 µL |

Bring volume to 500 mL with dd$H_2O$ and sterile filter.
Add NP-40 and protease inhibitors right before use.

**Reverse Crosslinking solution**

| Stock | Final | For 0.9 mL (for 6 samples) |
|---|---|---|
| 1X TE | | 720 µL |
| 20 mg/ml Proteinase K, 20X | 1X | 45 µL |
| 10% SDS | 1% | 90 µL |
| 10 mg/ml RNaseA (100X) | 1X | 9 µL |
| 5 M NaCl | 1X | 36 uL |

**Orange G 6X gel loading dye**

| Stock | Final | For 50 mL |
|---|---|---|
| Orange G | 0.1% w/v | 500 mg |
| 1 M Tris-HCl pH 7.5 | 10 mM | 500 µL |
| 0.5 M EDTA pH 8.0 | 60 mM | 6 mL |
| Glycerol | 60% | 30 mL |

Bring volume to 50 mL with dd$H_2O$ and sterile filter. Orange G will migrate @ ~50 bp in a 1% agarose gel.

**Materials and Reagents**

| Item | Manufacturer / Supplier | Catalog # |
|---|---|---|
| Low-retention pipet tips | VWR | 76322 |
| NP-40 Alternative | Millipore Sigma | 492018 |
| Protease Inhibitor Cocktail (EDTA-free) | Sigma-Aldrich | 5056489001 |
| MNase (micrococcal nuclease) | Worthington Biochem | LS004798 |
| EGTA | bioWORLD | 40520008 |
| Proteinase K | Viagen Biotech | 501-PK |
| RNase A, DNase- and proteinase-free | ThermoFisher | EN0531 |
| SDS (sodium dodecyl sulfate) | Sigma-Aldrich | L3771 |
| Tris-EDTA buffer | Sigma-Aldrich | 93283 |
| Phenol:Chloroform:Isoamylic alcohol (PCI) | Sigma-Aldrich | P2069 |
| 5PRIME Phase Lock Gel Light tubes | Quantabio | 2302820 |
| Low-retention tubes (DNA LoBind) | Eppendorf | 0030108418 |
| DNA Clean & Concentrator kit | Zymo Research | D4034 |
| Agarose | VWR | 97062 |
